# Quantitative profiling of JMJD6-catalysed lysine hydroxylation reveals a graded, residue-dependent readout of oxygen availability

**DOI:** 10.64898/2026.06.08.730680

**Authors:** Pallavi Kesavan, Sophie M. Stuermer, Katrin Räbel, Oliver Popp, Philipp Mertins, Matthew E. Cockman, Yoichiro Sugimoto

## Abstract

Enzyme-catalysed protein hydroxylation links cellular oxygen availability to protein function, as exemplified by the hydroxylation of hypoxia-inducible factor (HIF). Whereas HIF hydroxylation occurs at three specific residues, lysine hydroxylation catalysed by Jumonji domain-containing protein 6 (JMJD6) occurs at many residues within lysine-rich regions across the proteome. How this widespread modification signals oxygen availability has remained incompletely understood, in part because hydroxylation within lysine-rich regions is difficult to detect. Here we developed an analytical workflow for comprehensive, accurate, and quantitative analysis of lysine hydroxylation in proteomic data from lysine-derivatised samples: optimised database searching improved peptide coverage across lysine-rich regions, while diagnostic immonium ion filtering increased the accuracy of hydroxylysine residue assignment. We further showed that stoichiometry estimated from peptide precursor ion intensities enables reliable comparison across residues and conditions. Applied to bromodomain (BRD) proteins, epigenetic readers containing lysine-rich regions extensively hydroxylated by JMJD6, the workflow revealed marked heterogeneity in the apparent kinetics of hydroxylation among target lysines, with interdependence between neighbouring hydroxylation events. Hypoxia suppressed hydroxylation progressively with increasing severity, preferentially affecting residues with slower apparent kinetics. Together, these findings demonstrate that lysine-rich regions can encode a graded, residue-dependent readout of oxygen availability, providing a framework for investigating how cells calibrate their responses to changes in oxygen levels.

## Introduction

Protein hydroxylation serves as a regulatory mechanism linking cellular oxygen availability to protein function^1^. For instance, prolyl hydroxylase domain (PHD) enzymes hydroxylate HIFα, thereby controlling HIFα stability and mediating oxygen-dependent transcriptional reprogramming^2^. PHDs belong to the 2-oxoglutarate-dependent dioxygenase (2-OGDD) superfamily, whose members catalyse hydroxylation reactions using molecular oxygen and 2-oxoglutarate as co-substrates^3^. Whereas PHD-mediated hydroxylation is restricted to a small set of target residues^2,4^, JMJD6 – another 2-OGDD – hydroxylates many more residues. It catalyses hydroxylation at the C5 position of at least 150 lysine residues across 48 substrates, with these residues frequently clustered within lysine-rich regions^5^. A periodate-based proteomic approach that selectively targets C5-hydroxylysine suggests that this modification is even more widespread^6^. The breadth of JMJD6 targets, together with the functional importance of lysine-rich regions^7^, raises the possibility that lysine hydroxylation influences multiple cellular processes in an oxygen-dependent manner. However, how lysine hydroxylation by JMJD6 varies across individual residues and responds to hypoxia has remained largely unresolved, in part because peptides from these regions are poorly recovered by standard mass spectrometry workflows and hydroxylation within them is difficult to measure accurately.

Lysine-rich regions form regulatory interfaces for molecular interaction and post-translational modification. The positively charged lysine side chain participates in electrostatic interactions and hydrogen bonding, and lysine is frequently enriched within intrinsically disordered regions that mediate multivalent interactions with RNA and other molecules. These properties allow lysine-rich regions to contribute to the assembly of biomolecular condensates^7^. Furthermore, lysine-rich regions are subject to a wide range of post-translational modifications (PTMs)^8^, including hydroxylation, methylation, acetylation, and ubiquitination, enabling dynamic regulation of protein interactions, localisation, and function. For example, modification of lysine-rich histone tails plays a central role in transcriptional control^9^. Among these PTMs, JMJD6-catalysed C5 hydroxylation is chemically distinct from the other modifications, which target the ε-amino group. C5 hydroxylation leaves the ε-amino group and its positive charge intact (Fig. 1a), while introducing a small polar group that adds new hydrogen-bonding capacity and may alter local side-chain interactions. As JMJD6 preferentially hydroxylates lysine-rich regions, this modification has the potential to regulate molecular interactions, condensate behaviour, and substrate recognition by lysine-modifying enzymes in an oxygen-dependent manner.

**Fig. 1.**
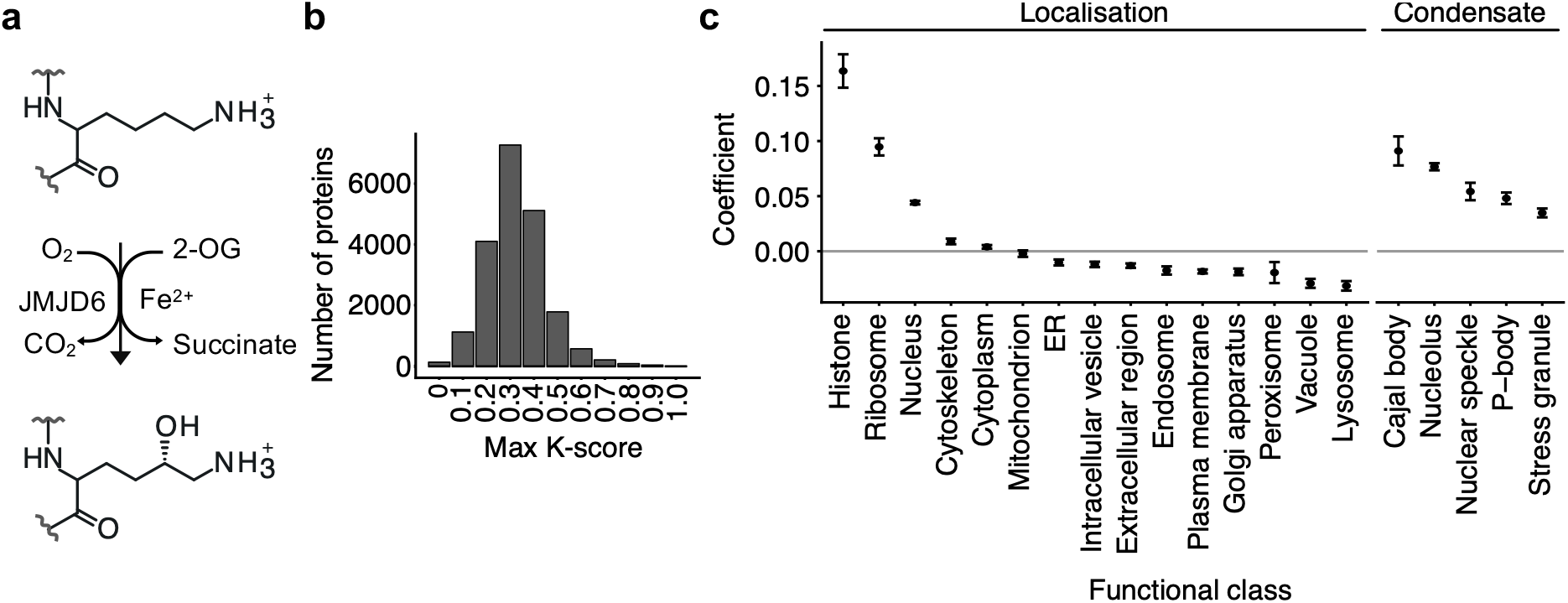
Lysine-rich regions are widespread and linked to nucleic acid binding and biomolecular condensates. **a.** Schematic of lysine C5 hydroxylation catalysed by JMJD6. **b.** Number of human proteins as a function of maximum K-score. **c.** Enrichment of lysine-rich regions among protein functional classes, grouped by subcellular localisation and condensate type. Data are presented as coefficients from a generalised additive model predicting maximum K-score (mean ± s.e.m.). The model accounts for the association between protein length and maximum K-score (see Supplementary Fig. 2e).

Bottom-up proteomics using liquid chromatography-tandem mass spectrometry (LC-MS/MS) is the primary method for studying protein hydroxylation^5,10^. In most workflows, proteins are digested with trypsin, which specifically cleaves polypeptide chains at the C-terminus of lysine and arginine residues, to generate peptides of suitable length and charge for efficient ionisation and identification. However, in lysine-rich regions, tryptic digestion produces peptides that are often too small to enable confident protein assignment. Although alternative proteases with different cleavage specificities could, in principle, address this limitation, they generally exhibit lower efficiency than trypsin and frequently generate peptides with suboptimal properties for LC-MS/MS analysis, such as high charge states^11^. Chemical derivatisation of lysine residues using propionic anhydride mitigates these issues by preventing tryptic cleavage at derivatised lysines and neutralising their charge^12,13^ (Supplementary Fig. 1). Although this approach enables recovery of MS-amenable peptides from lysine-rich regions, it introduces analytical complexity through incomplete derivatisation and altered digestion patterns. Accurate detection of hydroxylation by LC-MS/MS is also intrinsically challenging. Spontaneous oxidation of residues such as methionine leads to a mass shift isobaric with lysine hydroxylation, which – together with the difficulty of localising the modification to specific residues – can result in false-positive identifications^4,14^. To date, the performance of existing data analysis strategies for studying hydroxylation in lysine-rich regions has not been systematically evaluated.

To address these challenges, we refined key steps of the LC-MS/MS-based analysis of lysine-derivatised samples. Optimisation of database search settings increased the recovery of peptides in lysine-rich regions. In addition, filtering peptides based on diagnostic immonium ions characteristic of hydroxylysine markedly improved confidence in hydroxylysine assignment. We further demonstrated that stoichiometry, calculated from peptide precursor ion intensities, as assigned by database search software, enables quantitative comparison of lysine hydroxylation across residues and conditions. To facilitate this analysis, we developed an R package, *ptm.stoichiometry*, which enables direct calculation of PTM stoichiometry from the output of the widely used MS search software MaxQuant^15^. Overall, our workflow supports higher-confidence assignment and comparative quantification of lysine hydroxylation in lysine-rich regions.

We applied this workflow to characterise lysine hydroxylation in BRD epigenetic reader proteins, which are extensively hydroxylated by JMJD6^5^. The apparent kinetics of hydroxylation varied substantially between lysine residues, and hydroxylation events at neighbouring residues showed interdependence. Across quantified residues, JMJD6-dependent lysine hydroxylation decreased in a graded manner with increasing severity of hypoxia. Oxygen sensitivity differed markedly between residues, with the largest effects at residues exhibiting slower apparent kinetics. Together, these findings reveal that JMJD6-catalysed lysine hydroxylation can encode oxygen availability as a graded, residue-dependent modification pattern, and provide a basis for investigating how this sensitivity contributes to cellular responses to oxygen availability.

## Results

### Lysine-rich regions are widespread and linked to nucleic acid binding and biomolecular condensates

To characterise lysine-rich regions, we employed the previously defined K-score^5^, a measure of local lysine enrichment calculated as the proportion of residues that are lysine within a sliding 10-amino-acid window. As a reference, histone tails – canonical lysine-rich regions^16^ – exhibited a median K-score of 0.3 (Supplementary Fig. 2a). Across the human proteome, 74% of proteins contained at least one region with K-score comparable to or higher than that of histone tails (K-score ≥ 0.3, Fig. 1b), indicating that lysine-rich sequences are prevalent. These regions shared physicochemical and functional characteristics. Regions with higher K-scores exhibited greater positive charge and were predicted to be intrinsically disordered (Supplementary Fig. 2b–d). Lysine-rich regions were enriched in proteins that bind nucleic acids and form RNA-associated biomolecular condensates^17^, including Cajal bodies, nucleoli, and nuclear speckles (Fig. 1c). These associations are consistent with positive charge promoting nucleic acid binding and intrinsic disorder facilitating multivalent interactions and condensate formation. Together, these findings suggest that lysine-rich regions are widespread and functionally relevant sequence features. However, little is known about how PTMs are distributed across the many closely spaced lysines in these regions, or how a modification such as hydroxylation varies in stoichiometry at individual residues and responds to physiological cues such as oxygen availability. This uncertainty partly reflects the difficulty of analysing dense lysine sequences, which motivates the approaches developed here.

### Optimising peptide recovery from lysine-rich regions

Lysine derivatisation of proteins, followed by tryptic digestion, enables the generation of peptides suitable for LC-MS/MS analysis by neutralising the charge of lysine residues and blocking tryptic cleavage at them. However, incomplete derivatisation generates heterogeneous peptide populations with variable cleavage patterns that fall outside the fixed cleavage rules assumed by standard database searches, making the corresponding hydroxylated peptides difficult to recover during data analysis. To optimise peptide recovery, we reanalysed a dataset from our previous study, in which JMJD6 target proteins were enriched, derivatised with propionic anhydride, digested with trypsin, and analysed by LC-MS/MS^5^. Because this dataset was enriched for lysine-rich proteins and hydroxylated lysines, it was well suited to workflow optimisation.

The completeness of lysine derivatisation determines how tryptic cleavage should be modelled during database searching. Previous studies of histone modifications typically assumed complete derivatisation^12,13^, in which all lysines are blocked, and cleavage is therefore restricted to arginine. In contrast, our earlier work on JMJD6-dependent hydroxylation assumed incomplete derivatisation – in which underivatised lysines remain available for tryptic cleavage – and allowed up to five missed cleavages in tryptic digests^5^. We therefore varied two coupled *in silico* digestion parameters: (i) whether cleavage was restricted to arginine or also permitted at lysine and (ii) the number of allowed missed cleavages. Propionylation, hydroxylation, and their combination were searched as variable modifications. Because lysine propionylation blocks tryptic cleavage, each propionylated residue contributes both a variable modification and a missed cleavage during database searching. Accordingly, the maximum number of variable modifications was matched to the maximum number of allowed missed cleavages per peptide. Database searches were performed using MaxQuant with a 1% FDR.

Allowing additional missed cleavages increased the number of peptide-spectrum matches (PSMs), with saturation at approximately seven missed cleavages (Fig. 2a). By contrast, restricting cleavage to arginine residues yielded the fewest PSMs, consistent with incomplete lysine derivatisation in highly lysine-rich regions. Notably, increasing the number of missed cleavages did not increase PSM counts in non-derivatised datasets, indicating that this effect is specific to derivatised samples rather than a generic search artefact. We therefore compared settings allowing two (standard workflows), five (previous study^5^), and seven (more permissive) missed cleavages. Permitting up to seven missed cleavages improved peptide identification within these regions (Fig. 2b). Consistent with this global trend, peptide coverage within lysine-rich regions of BRD2–4 – previously shown to be extensively hydroxylated by JMJD6^5^ – was enhanced under more permissive settings (Supplementary Fig. 3). Together, these results establish search settings that recover lysine-rich peptides efficiently.

**Fig. 2.**
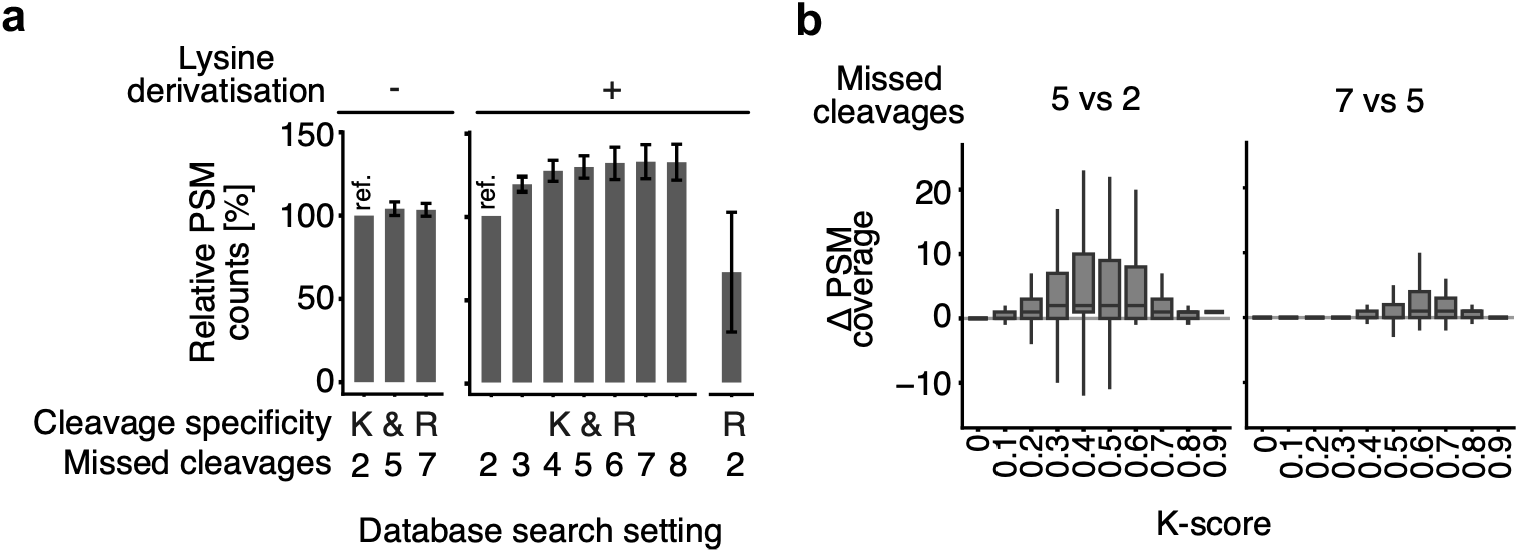
Optimised database search settings improve peptide recovery from lysine-rich regions. **a.** Bar charts showing the effect of varying *in silico* digestion parameters on peptide-spectrum match (PSM) counts. Data are normalised to the reference setting (ref.) defined as: cleavage at lysine (K) and arginine (R) with two missed cleavages. Bars show mean ± s.d. Data from BRD protein-enriched samples: left, immunoprecipitation without derivatisation; right, JQ1 enrichment with derivatisation. The maximum number of allowed missed cleavages was matched to the maximum number of variable modifications per peptide, except when cleavage was restricted to arginine, in which case two missed cleavages and seven variable modifications were allowed. **b.** Box plots showing differences in per-residue PSM coverage (Δ PSM coverage) stratified by K-score, for searches allowing 5 versus 2, and 7 versus 5 missed cleavages.

### Filtering peptides with diagnostic immonium ions improves the accuracy of hydroxylysine identification

We next sought to improve the accuracy of hydroxylysine assignment from MS data analysed at 1% FDR. We first asked whether the lysine residues assigned as hydroxylated by database searching contained a substantial fraction of false positives, using two orthogonal criteria. First, since JMJD6 is the lysine hydroxylase with the broadest substrate repertoire^5^, its inactivation should substantially reduce lysine hydroxylation levels. However, JMJD6 inactivation had only a modest effect on the number of lysine residues assigned as hydroxylated, many of which were detected in both JMJD6-inactivated and wild-type cells (Fig. 3a left panel and Supplementary Fig. 4a). Second, sequences surrounding residues assigned as hydroxylated were significantly enriched for methionine compared with those surrounding all lysines in the proteome (Fig. 3a left panel, two-sided Fisher’s exact test, *P* < 10^−10^). Since artefactual methionine oxidation is prevalent in proteomic datasets and produces the same mass shift as lysine hydroxylation, enrichment of methionine near residues assigned as hydroxylysine is consistent with spectral misassignment. Collectively, these findings suggest widespread misassignment of lysine hydroxylation, which is at least in part attributable to methionine oxidation.

**Fig. 3.**
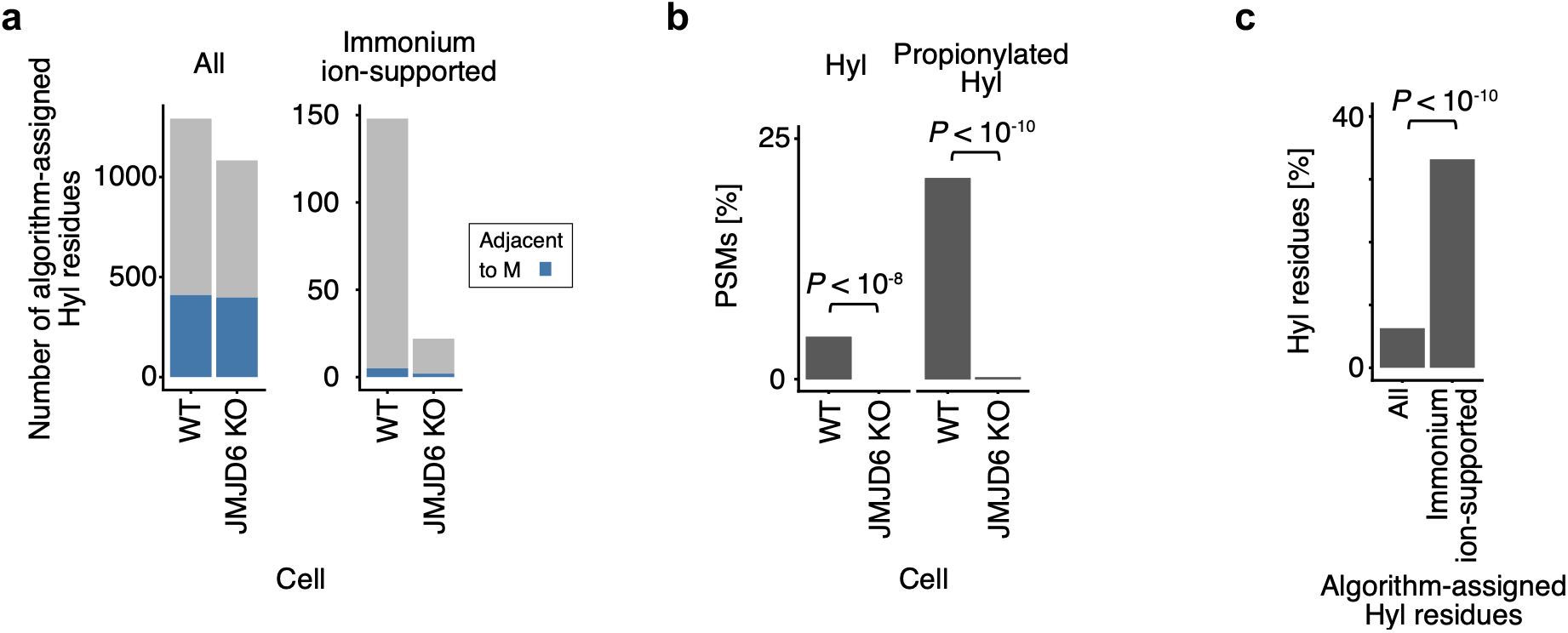
Diagnostic immonium ion filtering improves the accuracy of hydroxylysine assignment. **a.** Stacked bar charts showing the number of lysine residues assigned as hydroxylated in wild-type HeLa cells (WT) or JMJD6-inactivated HeLa cells (JMJD6 KO); blue indicates the subset of assignments in which a methionine residue lies within two residues of the assigned hydroxylated lysine. The left panel shows data for all hydroxylysine (Hyl) residues assigned by the software; the right panel shows data for hydroxylated residues supported by diagnostic immonium ions. Only residues supported by at least one peptide-spectrum match (PSM) in both WT and JMJD6 KO datasets were analysed. **b.** Bar charts showing the proportion of PSMs containing diagnostic immonium ions among peptides spanning the lysine-rich region of BRD4 (residues 535–555) in WT and JMJD6 KO HeLa datasets. The left and right panels show data for the diagnostic immonium ion of Hyl and propionylated Hyl, respectively. Proportions were compared using a two-sided Fisher’s exact test. n = 752 and 586 PSMs for WT and JMJD6 KO, respectively. **c.** Bar charts showing the proportion of software-assigned hydroxylated lysine residues that were identified by manual inspection of spectra in our previous work^5^, among all software-assigned residues (left) and immonium ion-supported residues (right). The two sets were compared using a two-sided Fisher’s exact test (n = 1,902 and 235, respectively).

To distinguish hydroxylysine peptides more accurately, we explored the use of diagnostic immonium ions. Modified and unmodified lysine residues can form stable cyclic immonium ions by neutral loss of ammonia during peptide fragmentation in LC-MS/MS analysis. These immonium ions can serve as diagnostic markers for certain lysine PTMs^18–20^. Analysis of peptide spectra of the lysine-rich region of BRD4 revealed ions with m/z values matching cyclic immonium ions of hydroxylysine and propionylated hydroxylysine. We assessed the specificity of these diagnostic immonium ions using JMJD6-inactivated cells as a negative control. Both diagnostic immonium ions were observed significantly more frequently in spectra from cells expressing JMJD6 than in JMJD6-inactivated cells, supporting their specificity (Fig. 3b and Supplementary Fig. 4b–d, two-sided Fisher’s exact test, *P* < 10^−8^ and *P* < 10^−10^ for hydroxylysines and propionylated hydroxylysines, respectively). However, the sensitivity of diagnostic immonium ion analysis may be limited. Only 4% and 21% of spectra from peptides spanning this region in JMJD6-expressing cells contained the hydroxylysine and propionylated-hydroxylysine diagnostic immonium ions, respectively (Fig. 3b), even though most are expected to contain hydroxylysines, given the high stoichiometry of JMJD6-dependent hydroxylation in the region^5^.

We then incorporated these two immonium ions as diagnostic signatures into database searches by MaxQuant to support identification and scoring of peptides containing hydroxylysines. We note that diagnostic immonium ions provide evidence that a peptide carries a hydroxylysine but do not by themselves localise the modification; site assignment continued to rely on database searching, with immonium support increasing overall confidence. For brevity, we refer to residues assigned as hydroxylated from immonium ion-filtered peptides as immonium ion-supported residues. Of 1,902 lysines initially assigned as hydroxylated, 235 residues were supported by diagnostic immonium ions (Supplementary Table). The immonium ion-supported residues were significantly enriched for previously validated hydroxylated residues compared with the unfiltered set (Fig. 3c, two-sided Fisher’s exact test, *P* < 10^−10^), indicating that the filtering selects for genuine hydroxylated residues. The incomplete overlap between these residues and those identified in our previous work likely reflects differences in database search settings including the greater number of permitted missed cleavages, differences in database search algorithms (PEAKS^21^ versus MaxQuant), and the limited sensitivity of immonium ion filtering. Re-applying the same two false-positive assessment criteria to these immonium ion-supported residues confirmed the gain in accuracy. Inactivation of JMJD6 now markedly reduced the number of assigned lysine hydroxylation events (143 in JMJD6-expressing versus 20 in inactivated cells), consistent with its role as a major lysine hydroxylase. The immonium ion-supported residues also lacked the methionine enrichment in surrounding sequences seen among unfiltered assignments (Fig. 3a right panel).

Several newly identified hydroxylated lysine residues, supported by peptides containing diagnostic immonium ions, are adjacent to previously reported ones, including those in ARF-like GTPase 6-interacting protein 4 (ARL6IP4) and dyskerin pseudouridine synthase 1 (DKC1) (Supplementary Fig. 5). These residues lie in highly lysine-rich regions that JMJD6 preferentially hydroxylates, and hydroxylation was detected only in cells with intact JMJD6. These observations indicate that they are bona fide JMJD6 targets. In contrast, a small subset of immonium ion-supported residues were hydroxylated in both JMJD6-expressing and JMJD6-inactivated cells. These included proteins previously reported to be hydroxylated at lysines, such as collagen type V^22^, supporting their assignment as genuine JMJD6-independent modifications. Overall, these results show that diagnostic immonium ion filtering improves the accuracy of hydroxylysine assignment, yielding a higher-confidence set of residues for the quantitative and biological analyses that follow.

### Precursor ion intensity-derived stoichiometry enables quantitative comparison of lysine hydroxylation

Reliable stoichiometry estimates are important for comparing lysine hydroxylation across residues and conditions, as well as for assessing its functional significance. We previously estimated stoichiometry using peptide precursor ion intensities, as assigned by database search software, as proxies for peptide abundance^5^. However, this approach can be confounded by stochastic peptide identification and variation in ionisation efficiency, particularly following lysine derivatisation. We therefore evaluated whether this approach resolves dynamic changes at individual residues and whether the resulting values correlate with those obtained using an established quantification method.

We first assessed whether stoichiometry captures dynamic changes at individual residues. To this end, we performed a temporal analysis of lysine hydroxylation of BRD2–4 using JQ1-affinity enrichment in JMJD6-inducible HeLa cells, in which JMJD6 was inactivated by CRISPR/Cas9-mediated genome editing and re-expressed under the control of a doxycycline-inducible promoter. Analysis was restricted to hydroxylated residues supported by diagnostic immonium ions and residues assigned in our earlier work^5^. Consistent with previous observations^5^, negligible lysine hydroxylation was detected in the absence of JMJD6 induction. Upon re-expression, stoichiometry increased progressively with the duration of JMJD6 induction (Fig. 4a and b, Supplementary Fig. 6). Normalisation of stoichiometry values to those of wild-type HeLa cells at each lysine confirmed a time-dependent increase at individual residues. Differences between time points were resolvable up to 18 hours, after which hydroxylation patterns closely resembled those of wild-type HeLa cells (Fig. 4c and Supplementary Fig. 6). Accordingly, extended time points (18- and 24-hour induction) were combined for subsequent analyses. Together, these findings indicate that precursor ion intensity-derived stoichiometry reliably resolves time-dependent changes in lysine hydroxylation at single-residue resolution.

**Fig. 4.**
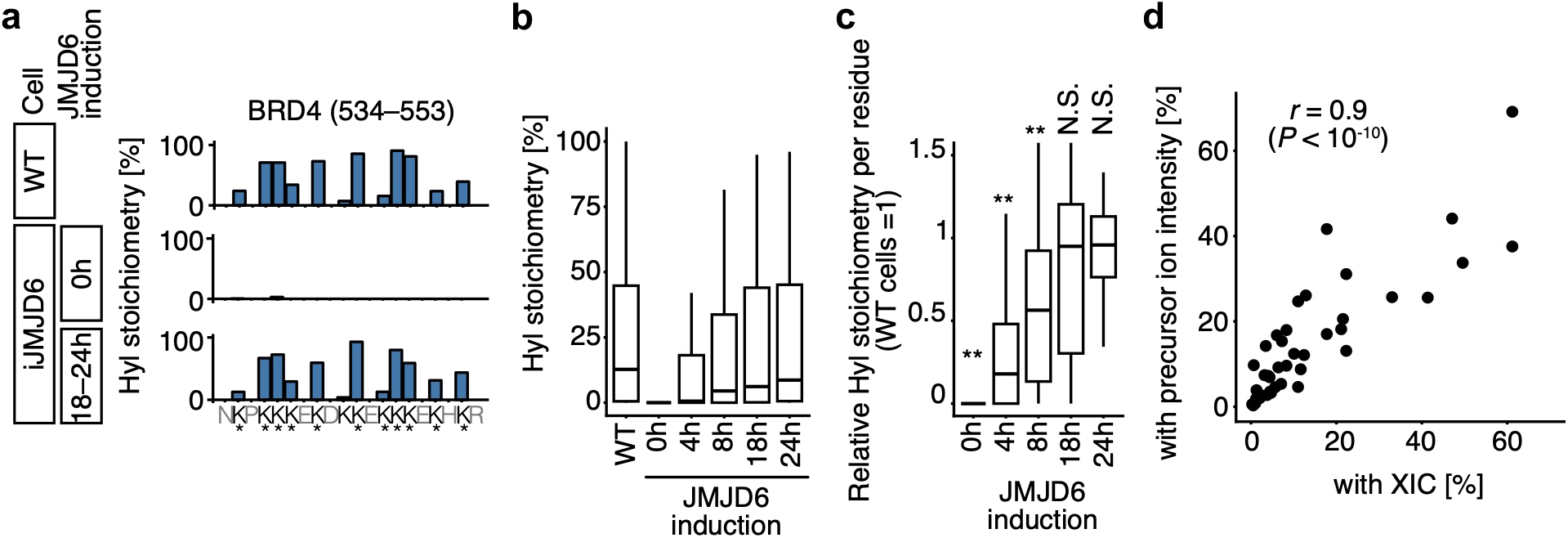
Peptide precursor ion intensity-derived stoichiometry enables quantitative comparison of lysine hydroxylation. **a.** Stoichiometry of hydroxylysine (Hyl) in the lysine-rich region of BRD4. Data for wild-type HeLa cells (WT) are compared with those for JMJD6-inducible (iJMJD6) HeLa cells with or without 18–24h of JMJD6 re-expression. Asterisks indicate residues with hydroxylation assignments supported by diagnostic immonium ions or assigned in our previous work^5^. **b.** Changes in Hyl stoichiometry upon re-expression of JMJD6 in JMJD6-inducible HeLa cells for the indicated duration. Data for wild-type HeLa cells are also shown. **c.** As in **b**, but showing Hyl stoichiometry per residue, normalised to that of WT HeLa cells. Data from each time point were compared with WT using a two-sided *t*-test. *P* values were adjusted for multiple comparisons using Holm’s method. \**P* < 0.05, \*\**P* < 0.005. N.S., not significant. **d.** Scatter plot comparing BRD2–4 hydroxylysine stoichiometry calculated by manual integration of XICs from non-derivatised samples with stoichiometry calculated from software-assigned peptide precursor ion intensities in derivatised samples. Each data point represents a residue–oxygen level–time point combination, with independent duplicate measurements for the 18–24h time point in derivatised samples (n = 48; see Supplementary Fig. 7b for individual data points). For **b** and **c**: n = 42.

We next assessed whether stoichiometries, obtained with our pipeline from derivatised samples, correlate with those from an established method — manual integration of extracted ion chromatograms (XICs) of non-derivatised samples. This analysis was restricted to hydroxylated residues reliably identifiable in non-derivatised samples; stoichiometries were compared across three induction time points and three oxygen levels (Supplementary Fig. 7a). The two approaches showed a strong correlation (Pearson’s *r* = 0.9, *P* < 10^−10^; Fig. 4d and Supplementary Fig. 7b), supporting the use of precursor ion intensity-derived stoichiometry from derivatised samples for comparative quantification across residues and conditions.

### Interdependent hydroxylation of neighbouring lysines contributes to residue-specific stoichiometry

Having optimised analytical parameters and established the suitability of the approach for comparative quantification, we applied this workflow to examine the apparent kinetics and oxygen sensitivity of JMJD6-dependent lysine hydroxylation in BRD2–4. These proteins were selected because they are epigenetic readers that play key roles in physiology and pathology^23^ and contain highly lysine-rich regions, in which many lysines are hydroxylated by JMJD6^5^.

A characteristic feature of JMJD6-mediated lysine hydroxylation is that modification often occurs at multiple lysines within a lysine-rich region, with variable stoichiometry. To investigate residue-dependent differences in stoichiometry, we analysed the apparent kinetics of lysine hydroxylation at individual residues. Specifically, we estimated the time required to reach half-maximal stoichiometry, hereafter referred to as *t_50_*, following JMJD6 induction under normoxia, using the time course data described above and isotonic regression. We note that *t_50_* is an apparent, rather than intrinsic, kinetic parameter for two reasons: (i) doxycycline treatment precedes JMJD6 protein induction by an appreciable lag and (ii) hydroxylation at different lysine residues may be interdependent. Nevertheless, the metric provides a useful basis for comparing apparent hydroxylation kinetics across different residues in cells. The *t_50_* varied substantially across residues. Residues with lower *t_50_* reached higher steady-state stoichiometry than those with higher *t_50_*, indicating that faster apparent kinetics are associated with higher hydroxylation stoichiometry (Fig. 5a and Supplementary Fig. 6).

**Fig. 5.**
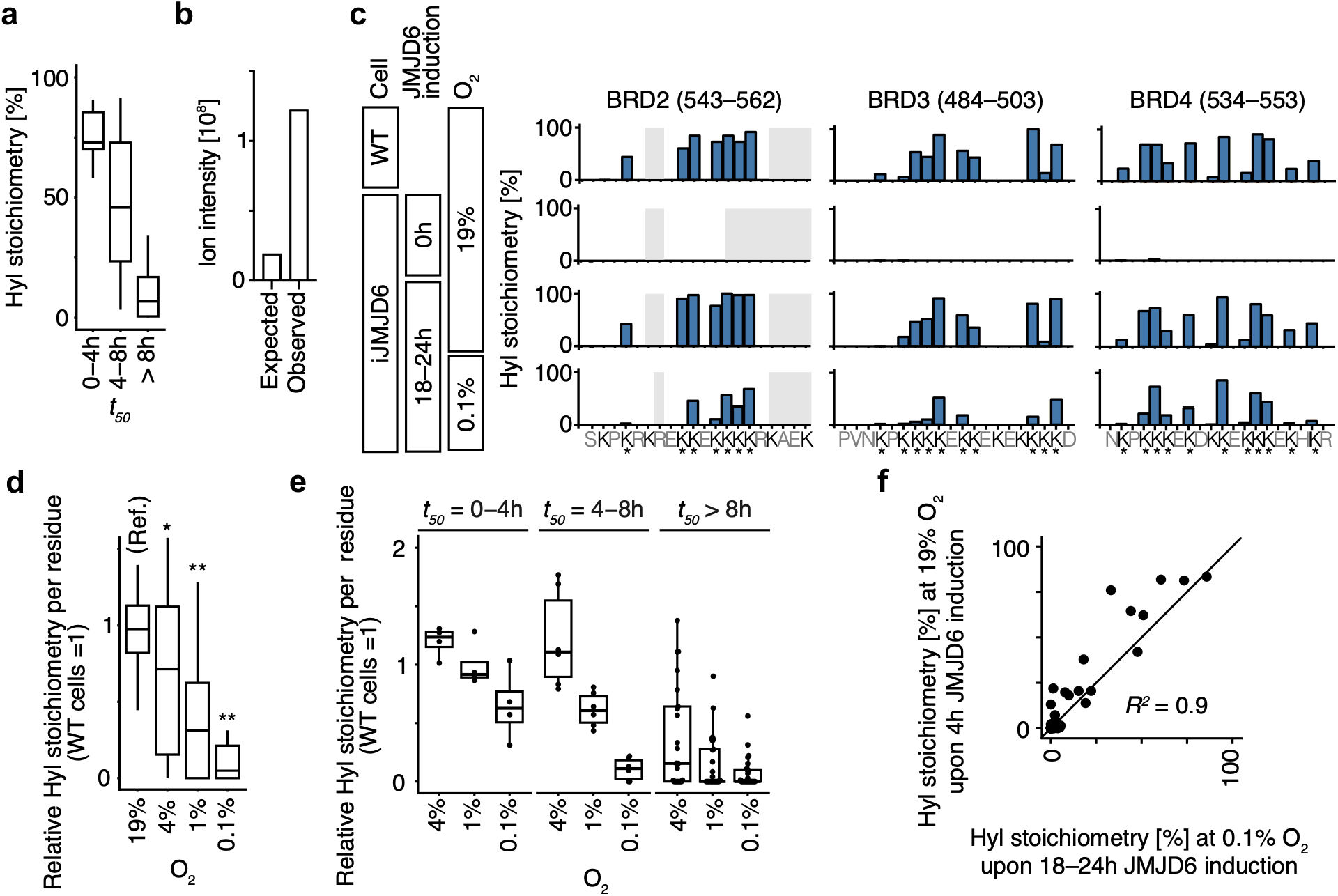
Hypoxia inhibits lysine hydroxylation in a residue-dependent manner. **a.** Hydroxylysine (Hyl) stoichiometry in wild-type (WT) HeLa cells as a function of the time required to reach half-maximal stoichiometry (*t_50_*) under normoxia. n = 9, 13, and 26 for *t_50_* = 0–4h, 4–8h, and > 8h, respectively. **b.** Bar plot comparing the observed XIC peak area of the doubly hydroxylated BRD4 (520–537) peptide with the value expected under a model assuming independent hydroxylation of K535 and K537. **c.** Hyl stoichiometry in the lysine-rich regions of BRD2–4. WT HeLa cells were compared with JMJD6-inducible (iJMJD6) HeLa cells with or without JMJD6 re-expression at 19% or 0.1% O_2_. Grey boxes indicate positions with insufficient data coverage (PSMs < 3). Asterisks indicate residues with hydroxylation assignments supported by diagnostic immonium ions or assigned in our previous work^5^. **d.** Hyl stoichiometry per residue in iJMJD6 HeLa cells after 18–24h of JMJD6 induction under the indicated oxygen levels, normalised to Hyl stoichiometry in WT HeLa cells at 19% O_2_. Each oxygen level was compared with the 19% O_2_ reference (Ref.) using a two-sided paired *t-*test. *P* values were adjusted for multiple comparisons using Holm’s method. \**P* < 0.05, \*\**P* < 0.005. n = 48. **e.** As in **d**, but stratified by *t_50_* under normoxia. n = 4, 6, and 25 for *t_50_* = 0–4h, 4–8h, and > 8h, respectively. **f.** Scatter plot comparing Hyl stoichiometry in iJMJD6 HeLa cells after 4h of JMJD6 induction under normoxia (19% O_2_) versus 18–24h of JMJD6 induction with concurrent exposure to 0.1% O_2_.

Because JMJD6 often hydroxylates multiple lysines within the same lysine-rich region, we next investigated whether hydroxylation at neighbouring residues was interdependent. We analysed robustly quantified peptides containing pairs of hydroxylated residues assigned to different *t_50_* categories, and assessed whether hydroxylation at one residue was associated with altered hydroxylation stoichiometry at the neighbouring residue. Higher-*t_50_* residues showed significantly higher stoichiometry when quantified from peptides in which neighbouring lower-*t_50_* residues were hydroxylated than when they were unmodified (Supplementary Fig. 8; paired two-sided t-test, *P* < 10^−5^). This pattern is consistent with coordinated hydroxylation at neighbouring lysine residues.

We further examined this relationship using a representative pair of residues, K535 and K537 of BRD4, for which hydroxylation was detectable by LC-MS/MS in non-derivatised samples. Under a model assuming independence between residues, the expected XIC peak area of the doubly hydroxylated peptide was calculated from the product of XIC peak areas of the singly hydroxylated forms, scaled by the XIC peak area of the unmodified peptide. The observed XIC peak area of the doubly hydroxylated species was 6.3-fold higher than expected under this model (Fig. 5b). This marked deviation indicates that hydroxylation at these residues does not occur independently. Together, these analyses demonstrate interdependence between hydroxylation events at neighbouring lysine residues, indicating that neighbouring modification states may contribute to the residue-specific apparent kinetics observed above.

### Hypoxia represses lysine hydroxylation in a residue-dependent manner

JMJD6 requires oxygen for catalysis^5,24,25^. Consistent with this requirement, we and others have shown, using distinct peptides that share residues 535–541 of the BRD4 lysine-rich region, that hydroxylation within this region is reduced under hypoxia both in cells^5^ and *in vitro*^25^. However, these analyses were restricted to individual peptides and reported only the total number of hydroxylation events per peptide. How hypoxia affects hydroxylation at individual residues and more broadly across BRD2–4, and whether it does so in a graded manner across physiological oxygen levels, remained unresolved.

To address these questions, we quantified hydroxylation stoichiometries in JMJD6-inducible HeLa cells subjected to concurrent JMJD6 induction and different oxygen levels for 18–24 hours. Lysine hydroxylation decreased in a graded manner with increasing hypoxic severity (Fig. 5c and d, Supplementary Fig. 9). JMJD6 abundance moderately increased under hypoxia, whereas BRD4 abundance was unaffected, suggesting that reduced hydroxylation reflects inhibition of JMJD6 catalytic activity rather than reduced enzyme or increased substrate abundance (Supplementary Fig. 10). Notably, inhibition of hydroxylation was evident at 4% O_2_ (Fig. 5d), indicating that JMJD6-catalysed lysine hydroxylation is sensitive to physiologically relevant changes in oxygen availability.

Oxygen sensitivity varied substantially across residues and was strongly associated with apparent hydroxylation kinetics. Residues with fast apparent kinetics (*t_50_* of 0–4 hours) were repressed only under severe hypoxia (0.1% O_2_), while they remained largely unaffected at more moderate hypoxia (1% and 4% O_2_). In contrast, residues with intermediate apparent kinetics (*t_50_* of 4–8 hours) showed graded repression across 0.1–4% O_2_. Residues with slow apparent kinetics (*t_50_* > 8 hours) were repressed at all tested hypoxic conditions (Fig. 5e). Thus, hypoxia inhibits lysine hydroxylation in a residue-dependent manner, with the greatest sensitivity observed at residues with slower apparent kinetics.

Interestingly, hydroxylation patterns observed under hypoxia resembled those seen at earlier time points of JMJD6 induction under normoxia. The stoichiometry profile after 18–24 hours of JMJD6 induction at 0.1% O_2_ most closely resembled the profiles observed after 4 hours of induction at normoxia or 4% O_2_, and after 8 hours at 1% O_2_ (Fig. 5f, Supplementary Fig. 11). These findings are consistent with the notion that hypoxia represses lysine hydroxylation by effectively slowing the apparent kinetics of JMJD6 activity according to the severity of hypoxia.

## Discussion

Lysine-rich regions present a persistent challenge for mass spectrometry-based proteomics. This difficulty may explain why the extensive substrate repertoire of JMJD6 has only recently become apparent^5,6^. Here, we developed an optimised workflow for the analysis of LC-MS/MS data from lysine-derivatised samples. Allowing multiple missed cleavages and variable modifications to account for incomplete lysine derivatisation by propionic anhydride increased peptide recovery from lysine-rich regions. Requiring diagnostic immonium ion support improved confidence in hydroxylysine assignments. Furthermore, we demonstrated that stoichiometry estimates derived from software-assigned peptide precursor ion intensities enabled comparative quantification of lysine hydroxylation across residues and conditions. Together, these advances establish a coherent workflow for analysing lysine hydroxylation within lysine-rich regions.

JMJD6-catalysed lysine hydroxylation is widespread and frequently occurs at multiple lysines within disordered lysine-rich regions^5^. This clustered, multi-site hydroxylation contrasts with that catalysed by factor-inhibiting hypoxia-inducible factor (FIH), another promiscuous 2-OGDD hydroxylase, which also modifies many substrates but does so at single asparaginyl or histidinyl residues within folded ankyrin repeat domains^10,26,27^. Analysis of BRD proteins revealed that apparent hydroxylation kinetics varied markedly across residues, even within the same lysine-rich region. These differences may reflect local sequence context and the interdependence we observed between neighbouring residues, which may arise from altered JMJD6 affinity or protein conformational changes after initial hydroxylation. Importantly, oxygen sensitivity also varied markedly across hydroxylated residues. Residues with slower apparent kinetics were more strongly repressed by hypoxia than those with faster kinetics, and hydroxylation patterns observed under hypoxia resembled those seen following short JMJD6 induction under normoxia. Collectively, these findings suggest that reduced oxygen availability slows the apparent accumulation of JMJD6-dependent lysine hydroxylation, resulting in residue-dependent repression of modification with increasing severity of hypoxia.

Residue-dependent oxygen sensitivity has important biological implications, potentially permitting progressive responses to increasing hypoxic severity. Notably, hydroxylation was already suppressed at 4% O_2_ and declined progressively down to 0.1% O_2_. This result indicates that JMJD6-catalysed lysine hydroxylation could provide a graded molecular readout of oxygen availability not only under severe hypoxia but also within the physiologically relevant range, given that most tissues are oxygenated at roughly 2–9% O_2_^28^. Individual substrates, such as BRD4, contain multiple hydroxylated residues with distinct oxygen sensitivities. Oxygen levels are therefore reflected in residue-specific patterns of hydroxylation within lysine-rich regions, which may shape protein function. More broadly, the same principle, applied across substrates, could enable fine-tuning of pathway activity in response to hypoxia. For instance, JMJD6 hydroxylates multiple splicing factors^5,24^; if these are hydroxylated with different oxygen sensitivities, pre-mRNA splicing could be adjusted progressively as oxygen levels decline. Finally, JMJD6 is overexpressed in several cancers, including neuroblastoma^29–31^, raising the possibility that increased JMJD6 abundance could alter apparent hydroxylation kinetics, potentially modulating the oxygen sensitivity of target proteins in pathophysiological contexts.

Although the workflow developed here improves confidence in hydroxylysine assignments and supports their comparative quantification, several limitations should be considered. While diagnostic immonium ions provide valuable evidence for the presence of hydroxylysine within a peptide, particularly in MS/MS spectra where backbone fragmentation is sparse, they cannot provide positional information. This matters because, in lysine-rich peptides, co-eluting hydroxylated positional isomers present a significant analytical challenge: co-fragmentation of peptides hydroxylated at different lysine residues can produce chimeric spectra, where overlapping site-determining ions may support conflicting localisation assignments. In addition, isobaric species may confound diagnostic-ion filtering. For example, propionylated hydroxylysine and lactyllysine exhibit identical mass shifts relative to unmodified lysine^8^, and thus the corresponding diagnostic immonium ions cannot be distinguished. Finally, although stoichiometry estimates derived from peptide precursor ion intensities are suitable for comparative analyses, absolute quantification requires synthetic peptide standards^32^. However, such standards are difficult to use at proteome-wide scale, particularly because hydroxylysine-containing synthetic peptides are not readily available.

Our biological analyses focused on BRD proteins, which contain well-defined JMJD6-targeted lysine-rich regions. Although the principles identified here – including residue-dependent apparent hydroxylation kinetics and oxygen sensitivity, as well as interdependence between neighbouring residues – are likely to apply to other JMJD6 targets, future studies extending this work across additional substrates will be needed to establish their generality. In addition, the roles of JMJD6^33^ and lysine-rich regions^7^ in health and disease suggest that JMJD6-mediated lysine hydroxylation has important functional consequences. However, establishing this will require studies that combine functional assays with measurement of lysine hydroxylation.

Taken together, this study provides a workflow that improves confidence in assigning lysine hydroxylation to individual residues within lysine-rich regions and supports comparative quantification across residues and conditions. The resulting analyses identify graded and residue-dependent oxygen sensitivity as a prominent feature of JMJD6-catalysed lysine hydroxylation in the BRD proteins examined. These findings provide a basis for investigating how this oxygen sensitivity contributes to adaptive cellular responses to changes in oxygen availability.

## Methods

### Data availability

LC-MS/MS data generated during this study are available from PRIDE under accession IDs: PXD079306, PXD079391, and PXD078942.

### Code availability

The R package developed during this study and the computational pipeline used for the data analysis are available on GitHub at https://github.com/YoichiroSugimoto/ptm.stoichiometry and https://github.com/YoichiroSugimoto/20241111_PTMs_in_lysine_rich_domains, respectively.

### Cell lines

JMJD6-inducible HeLa cells, in which JMJD6 was inactivated and re-expressed from a doxycycline-inducible promoter, were generated previously^5^. Cell identity was confirmed by STR profiling, and cells were confirmed to be free of mycoplasma contamination. HeLa cells were maintained in DMEM (Thermo Fisher Scientific, no. 31966047) supplemented with 10% tetracycline-free FBS (Pan Biotech, no. P30-3602) at 37 °C in 5% CO_2_. JMJD6-inducible HeLa cells were maintained under the same conditions with medium containing 1 µg/mL puromycin (InvivoGen, no. ant-pr-1). JMJD6 expression was induced by treatment with 400–600 ng/mL doxycycline (Sigma-Aldrich, no. D9891-1G). Hypoxic incubation was performed using a Whitley H35 HEPA Hypoxystation or a Baker Ruskinn InvivO_2_ 500 at the indicated oxygen levels, 5% CO_2_, and 37 °C.

### Antibodies

The following antibodies were used for immunoblotting analysis: anti-BRD4 (Abcam, ab128874) and anti-JMJD6 (Santa Cruz Biotechnology, sc-28348).

### Affinity purification of BRD proteins and sample preparation for MS analysis

Samples of BRD proteins for LC-MS/MS analysis were prepared as previously described^11^, with minor modifications. The procedure involves the following steps:

#### Preparation of JQ1-coated beads

Dynabeads MyOne Streptavidin C1 (10 µL; Thermo Fisher Scientific, no. 65001) was washed twice with native IP buffer (20 mM Tris-HCl, pH 7.5; 100 mM NaCl; 0.5% Igepal CA-630; 5 µM MgCl_2_). The beads were resuspended in 100 µL of the same buffer. JQ1-biotin (100 pmol; MedChemExpress, HY-145667) was added, and the samples were incubated at 4 °C for 1 hour with rotation to allow coupling. The beads were washed twice with native IP buffer to remove unbound JQ1-biotin and resuspended in 10 µL of native IP buffer.

#### Pulldown of BRD proteins

Cells were grown on 15-cm dishes, washed twice with ice-cold PBS, and lysed with 1 mL of RIPA buffer (Serva Electrophoresis, no. 3924402) supplemented with HALT protease inhibitor (Thermo Fisher Scientific, no. 78445) and 1 mM DMOG (Sigma-Aldrich, no. 40001-50MG). Genomic DNA was digested by adding Benzonase at 1:1,000 (v/v) (Sigma-Aldrich, no. E1014-25KU) to the lysate. The lysate was clarified by centrifugation at 14,000*g* for 5 min at 4 °C. JQ1-coated beads (10 µL) were added to 1 mL of lysate containing 5 mg total protein, and the mixture was incubated with rotation for 2 h at 4 °C. The beads were washed three times with RIPA buffer. JQ1-bound proteins were eluted and subjected to SDS-PAGE. Gel regions containing BRD2–4 were excised and cut into small pieces. In a subset of experiments, cells were treated with 500 µM DMOG prior to harvesting and, DMOG was omitted from the lysis buffer. Differences in conditions between datasets are summarised in Supplementary Table.

#### Lysine derivatisation

Gel pieces were washed overnight at 4 °C in 50% methanol and 5% acetic acid, then washed for an additional 5 min at room temperature. Gel pieces were dehydrated with acetonitrile, rehydrated in 80 µL of 100 mM ammonium bicarbonate, and treated with 20 µL of 25% propionic anhydride (Sigma-Aldrich, no. 240311-50G) in acetonitrile. The reaction pH was immediately adjusted to 8 by addition of 8 µL of ammonium hydroxide (Sigma-Aldrich, no. 5438300250). Samples were incubated at 51 °C for 20 min with shaking at 2,000 rpm. The supernatant was removed, the gel pieces were dehydrated with acetonitrile, and the propionylation reaction was repeated once.

#### Trypsinolysis

Gel pieces were rehydrated in 100 µL of 2.5 ng/µL Trypsin Platinum (Promega, no. VA9000) in 50 mM ammonium bicarbonate, and samples were incubated at 37 °C overnight with shaking at 1,200 rpm. Tryptic peptides were collected from residual digest buffer and then extracted sequentially with 50% and 80% acetonitrile, each containing 5% formic acid. Extracted peptides were purified using Evotip Pure tips (Evosep Biosystems, no. EV2013) according to the manufacturer’s instructions.

### Mass spectrometry

LC-MS/MS analyses were performed using two acquisition workflows. For experiments conducted at the Max Delbrück Center for Molecular Medicine, peptides were separated using an EASY-nLC 1200 system (Thermo Fisher Scientific) coupled online to an Orbitrap Exploris 480 mass spectrometer (Thermo Fisher Scientific). For each run, 2 µL of peptide solution in solvent A (3% acetonitrile, 0.1% formic acid) was injected. Samples were loaded onto a reversed-phase column at 1 µL/min and eluted at a constant flow rate of 250 nL/min using a 110 min gradient from 2% to 90% solvent B (90% acetonitrile, 0.1% formic acid). Full gradient profile: 2%–20% B over 67 min, 20%–30% B over 20 min, 30%–60% B over 10 min, ramping to 90% B for column washing, followed by re-equilibration.

The mass spectrometer was operated in data-dependent acquisition (DDA) mode. Full MS1 scans were acquired at a resolution of 60,000 (m/z 350–1800), followed by higher-energy collisional dissociation (HCD) fragmentation of the top precursors with a cycle time of 1 s. MS2 scans were acquired at a resolution of 15,000, with an isolation window of 1.3 m/z, and normalised collision energy of 28%. Dynamic exclusion was set to 20 s. Only precursor ions with charge states between 2+ and 6+ and intensities above 50,000 were selected for fragmentation. The maximum injection time for MS2 was 100 ms.

For experiments conducted at The Francis Crick Institute, samples were prepared and analysed by LC-MS/MS on an Orbitrap Eclipse Tribrid mass spectrometer (Thermo Fisher Scientific), using methods described previously^5^. The workflow used for each dataset is summarised in Supplementary Table.

### Overview of data analyses

Data analyses were performed in R (4.5.1), using the following packages: data.table (1.17.8), dplyr (1.1.4), stringr (1.5.2), and ggplot2 (4.0.0). Reference human protein sequences were downloaded from UniProt (UP000005640_9606). K-score, predicted disorderedness (using IUPred2A^34^), and local charge were calculated in the previous study^5^. In addition to data generated in this study, MS data from HeLa cells reported by Cockman *et al*.^5^ (PRIDE ID: PXD031221) were analysed.

### Analysis of the association between functional protein classes and maximum K-score

A generalised additive model was used to assess the association between functional protein classes and K-score. A model using a thin plate regression spline was constructed to predict the maximum K-score of a protein based on its functional class and length. The gam function of the mgcv package (1.9-1) was used for implementation. The coefficients for protein functional classes were then analysed.

### Diagnostic ion identification

To identify diagnostic immonium ions of hydroxylysine, public MS data^5^ generated from proteins extracted from HeLa cells with or without JMJD6 inactivation and enriched using JQ1-coated beads were analysed. FragPipe (22.0)^35^, MSFragger (4.1), IonQuant (1.10.27), diaTracer (1.1.5), and Philosopher (5.1.1) were used with the following parameters: search_enzyme_cut_1 = K, allowed_missed_cleavage_1 = 5, search_enzyme_cut_2 = R, allowed_missed_cleavage_2 = 2, variable_mod_01 = 15.9949 M 3, variable_mod_02 = 42.0106, variable_mod_10 = 15.9949 K 5, variable_mod_11 = 56.026215 K 5, variable_mod_12 = 72.021126 K 5, and max_variable_mods_per_peptide = 5.

Diagnostic immonium ions for hydroxylysine (100.0762 Da) and propionylated hydroxylysine (156.1025 Da) were confirmed in peptides covering the lysine-rich region of BRD4 (residues 535–555), and subsequently used to configure diagnostic immonium ion-aware database searches in MaxQuant.

### Database search

Database searches were performed using MaxQuant (2.6.2.0). The following parameters were used: fixed modifications: carbamidomethylation (C); variable modifications: Oxidation (M), Acetylation (Protein N-Terminus), Propionylation (C_3_H_4_O at K), Oxidation (O at K) and Oxidised Propionylation (C_3_H_4_O_2_ at K); diagnostic ions: Oxidation at K (C_5_H_9_ON) and Oxidised propionylation (C_8_H_13_O_2_N). Unless otherwise specified, database searches allowed up to seven missed cleavages and seven variable modifications per peptide.

To improve analysis speed, the initial search considered only oxidation (M), acetylation (protein N-term), and propionylation as variable modifications; the main search included all modifications listed above, with hydroxylysine and propionylated-hydroxylysine immonium ions as diagnostic markers.

### Hydroxylated lysine residues with immonium ion support

Hydroxylation assignments supported by peptides containing diagnostic immonium ions were pooled across all analysed datasets to generate a consensus list of immonium ion-supported residues.

### Calculation of the stoichiometry of PTMs

Stoichiometry was calculated as described previously^5^, using the following formula:

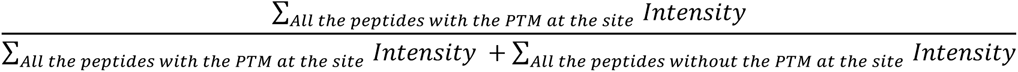

For comparisons between conditions, when multiple datasets were available for the same condition, the dataset with the highest coverage was selected as representative. Hydroxylation stoichiometry was calculated for lysines whose hydroxylation was identified by peptides with diagnostic immonium ions and/or by manual inspection of spectra in our previous work.

### Calculation of the time required to reach half-maximal stoichiometry (*t_50_*) upon JMJD6 induction

The time required to reach half-maximal stoichiometry (*t_50_*) was calculated using isotonic regression. Data from HeLa cells and JMJD6-inducible HeLa cells after 0, 4, 8, 18, and 24 hours of JMJD6 induction were used. For isotonic regression, genetically unmodified HeLa cells were assigned an induction time of 100 hours to represent prolonged JMJD6 expression, as the model requires finite numeric values.

## Author contributions

M.E.C. and Y.S. conceived the project. S.M.S., K.R., and O.P. performed experiments. P.K., S.M.S., M.E.C., and Y.S. analysed the data. P.K., S.M.S., O.P., P.M., M.E.C., and Y.S. contributed to the interpretation of the data. P.K., S.M.S., M.E.C., and Y.S. wrote the manuscript.

## Supplementary Table

Overview of the datasets and the stoichiometry of lysine hydroxylation calculated from each dataset.

## Appendix

Representative MS/MS spectra of non-derivatised peptides quantified by XIC analysis.

## Acknowledgements

We thank Sugimoto group members and Peter Ratcliffe (Francis Crick Institute) for support and discussion. This work was supported by the Francis Crick Institute which receives its core funding from Cancer Research UK (CC2092), the UK Medical Research Council (CC2092), and the Wellcome Trust (CC2092). We are grateful to Helen Flynn and Mark Skehel from the Proteomics STP at The Francis Crick Institute for their valued contributions to the work. For the purpose of Open Access, the author has applied a CC BY public copyright licence to any Author Accepted Manuscript version arising from this submission.

---

**Supplementary Fig. 1.**
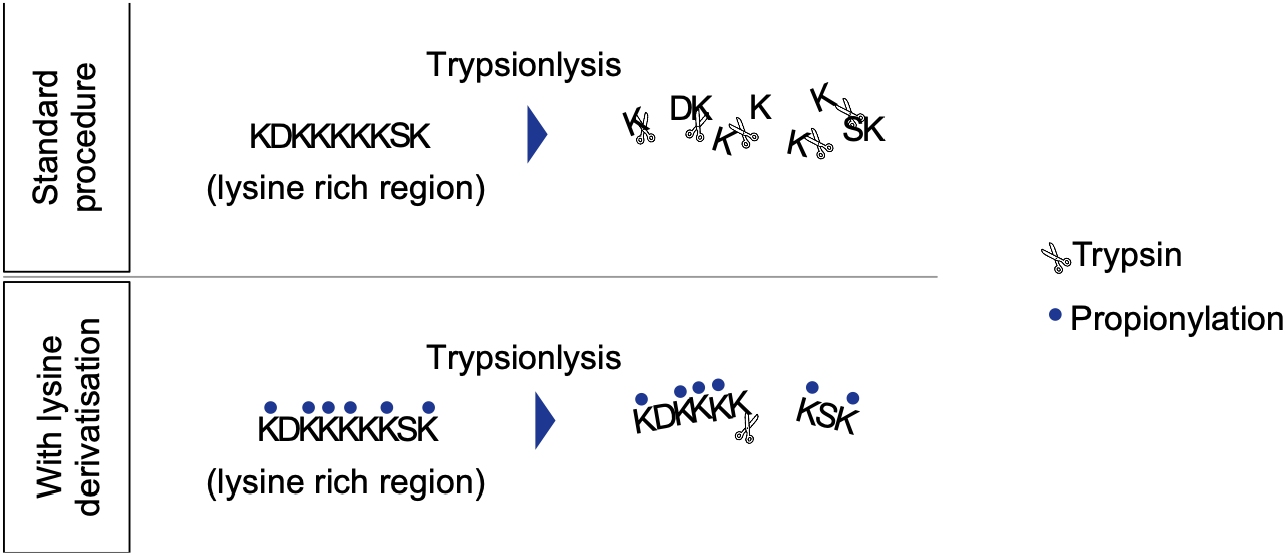
Schematic showing the effect of lysine derivatisation on trypsinolysis of lysine-rich regions.

**Supplementary Fig. 2.**
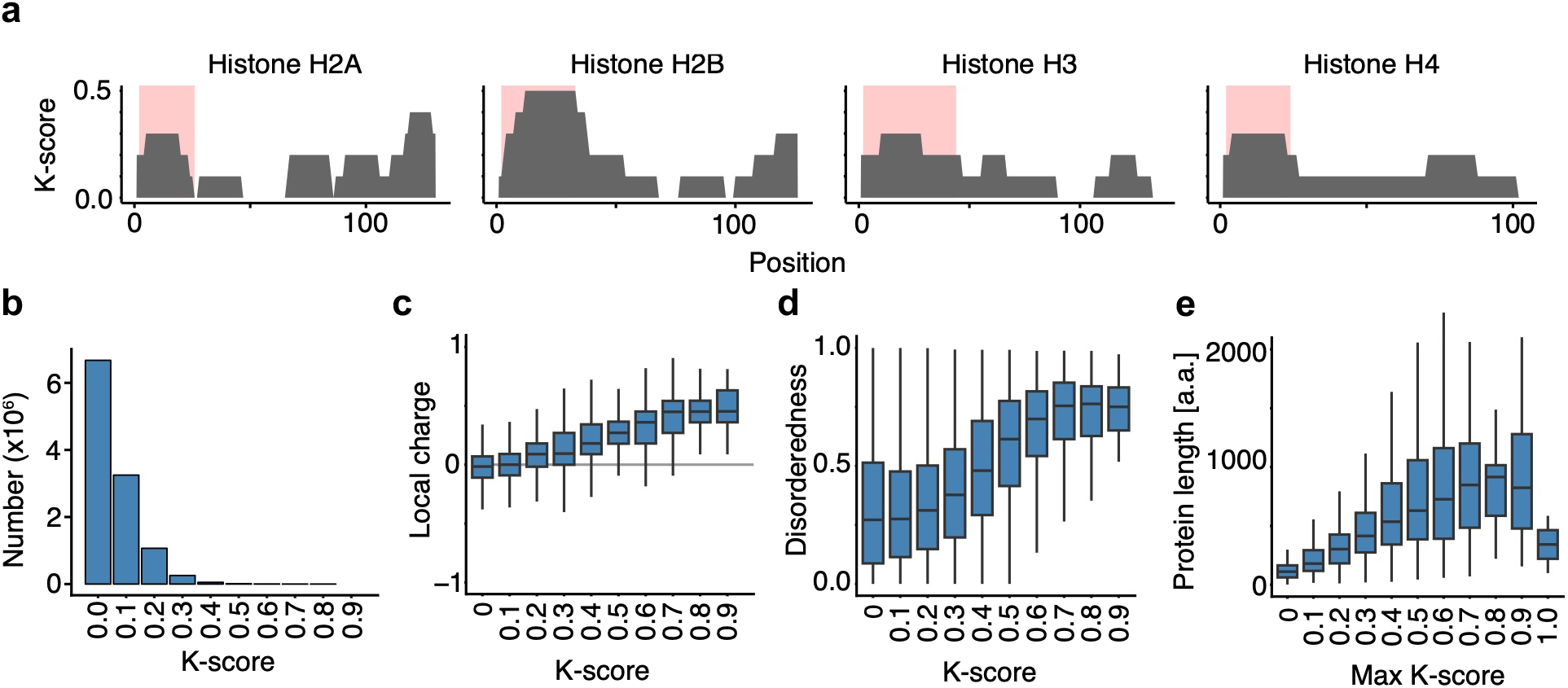
Characteristics of lysine-rich regions. **a.** Area plots showing K-score profiles across histone proteins. N-terminal histone tails are highlighted in red. **b.** Bar chart showing the distribution of K-scores calculated for each residue across the human proteome. **c.** Distribution of local charge as a function of K-score. Local charge was calculated over a window of ±5 residues. All sliding windows across the human proteome were analysed. **d.** As in **c**, but showing disorderedness as a function of K-score. Disorderedness was predicted using IUPred2A^34^. **e.** Distribution of protein length (amino acids; a.a.) as a function of maximum K-score.

**Supplementary Fig. 3.**
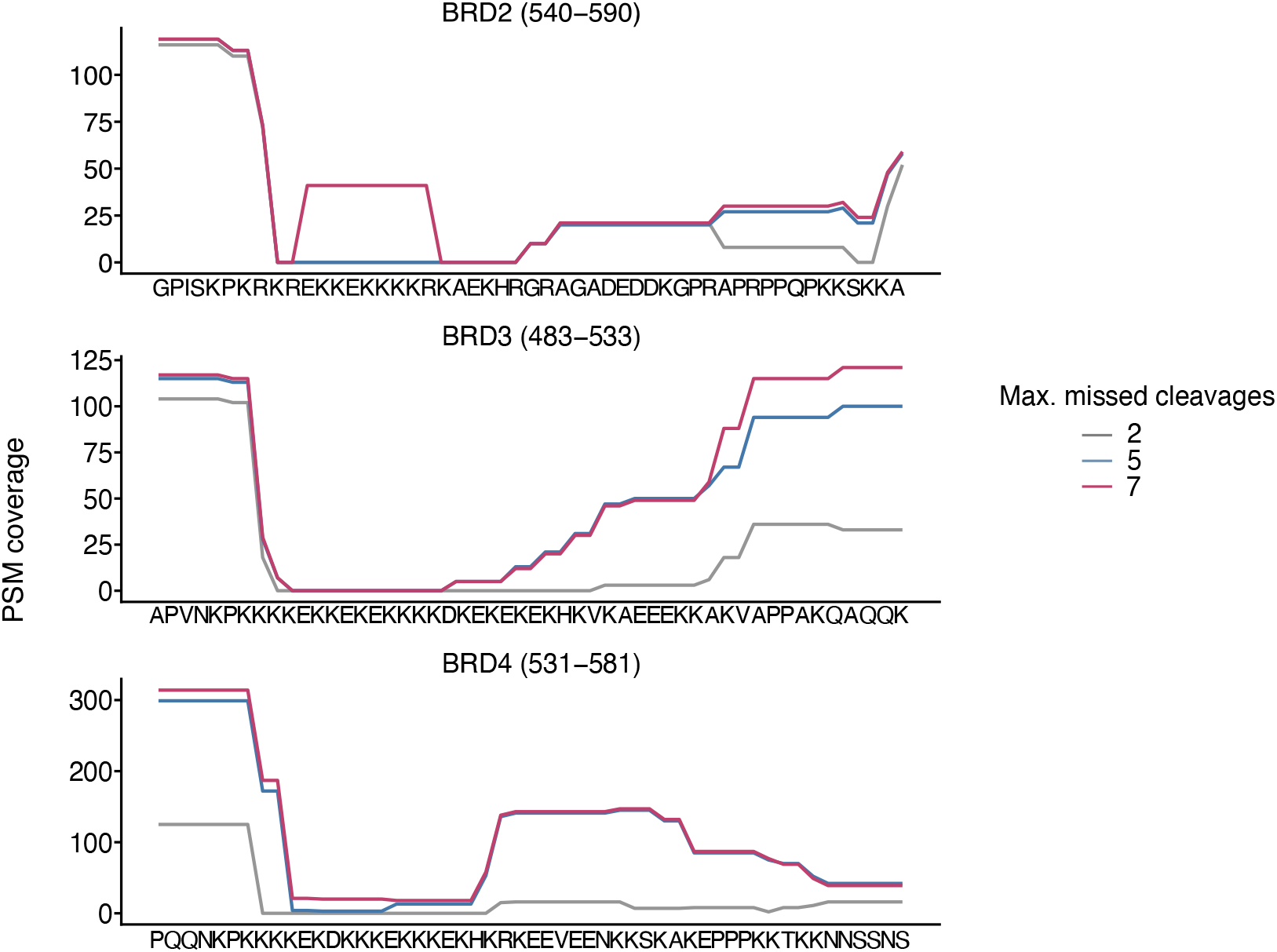
Optimised database search settings improve peptide recovery from lysine-rich regions of the BRD proteins. Peptide-spectrum match (PSM) coverage across lysine-rich regions of BRD2–4 under database search settings allowing 2, 5, or 7 missed cleavages. In each search, the maximum number of allowed missed cleavages was matched to the maximum number of variable modifications per peptide.

**Supplementary Fig. 4.**
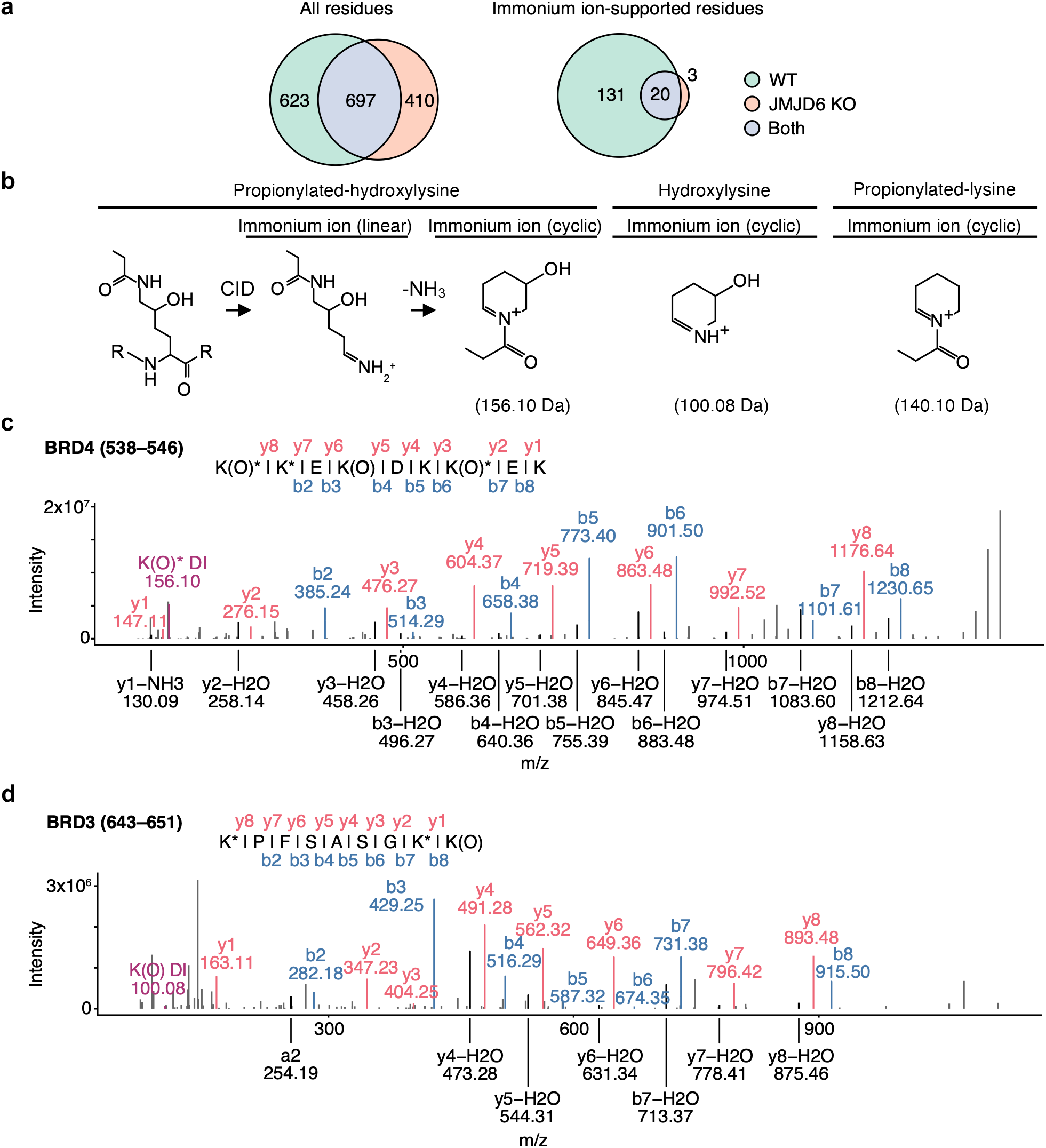
Diagnostic immonium ions of hydroxylysine and propionylated hydroxylysine. **a.** Number of lysine residues assigned as hydroxylated in wild-type (WT) and JMJD6-inactivated (JMJD6 KO) HeLa cells. Data are shown for all hydroxylysine assignments and for assignments supported by diagnostic immonium ions. **b.** Schematic of diagnostic immonium ions for propionylated hydroxylysine hydroxylysine, and propionylated lysine (shown for comparison). For propionylated hydroxylysine, the scheme shows collision-induced dissociation (CID) generating a linear immonium ion that loses ammonia (-NH_3_) to form the cyclic ion. **c.** MS/MS spectrum of a BRD4 peptide (residues 538–546), showing the diagnostic immonium ion peak for propionylated hydroxylysine. Assigned b- and y-ions are annotated Asterisks indicate propionylation, O indicates hydroxylation, and DI indicates the diagnostic immonium ion. **d.** As in **c**, but for a BRD3 peptide (residues 643–651), showing the diagnostic immonium ion peak for hydroxylysine.

**Supplementary Fig. 5.**
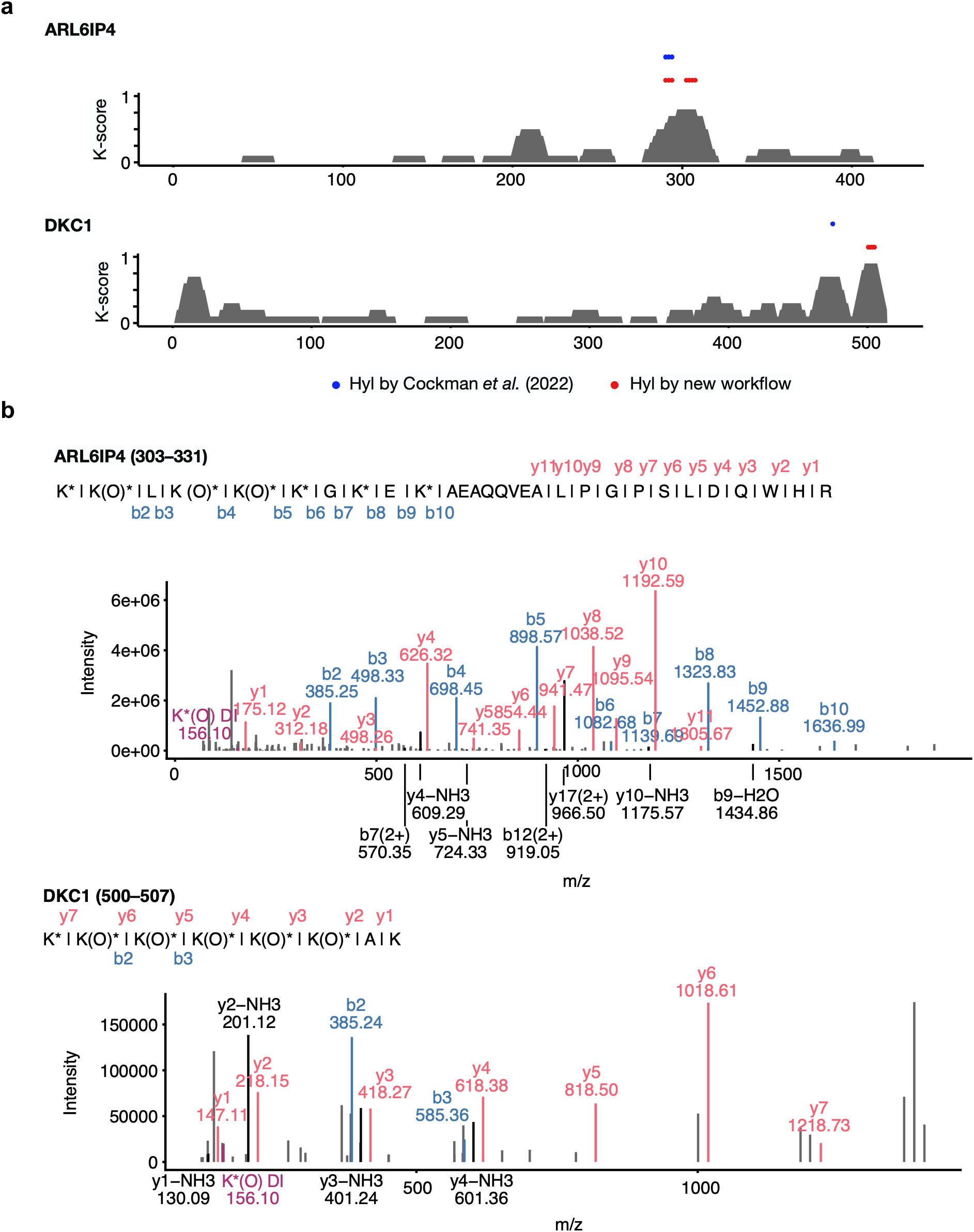
Newly identified JMJD6-dependent hydroxylated residues. **a.** K-score profiles across ARL6IP4 and DKC1. Lysine residues identified as hydroxylated in our previous work^5^ (blue) and newly identified in this study using diagnostic immonium ions (red) are indicated. **b.** MS/MS spectra of the ARL6IP4 peptide (residues 303–331) and the DKC1 peptide (residues 500–507). Assigned b- and y-ions are annotated. Asterisks indicate propionylation, O indicates hydroxylation, and DI indicates the diagnostic immonium ion.

**Supplementary Fig. 6.**
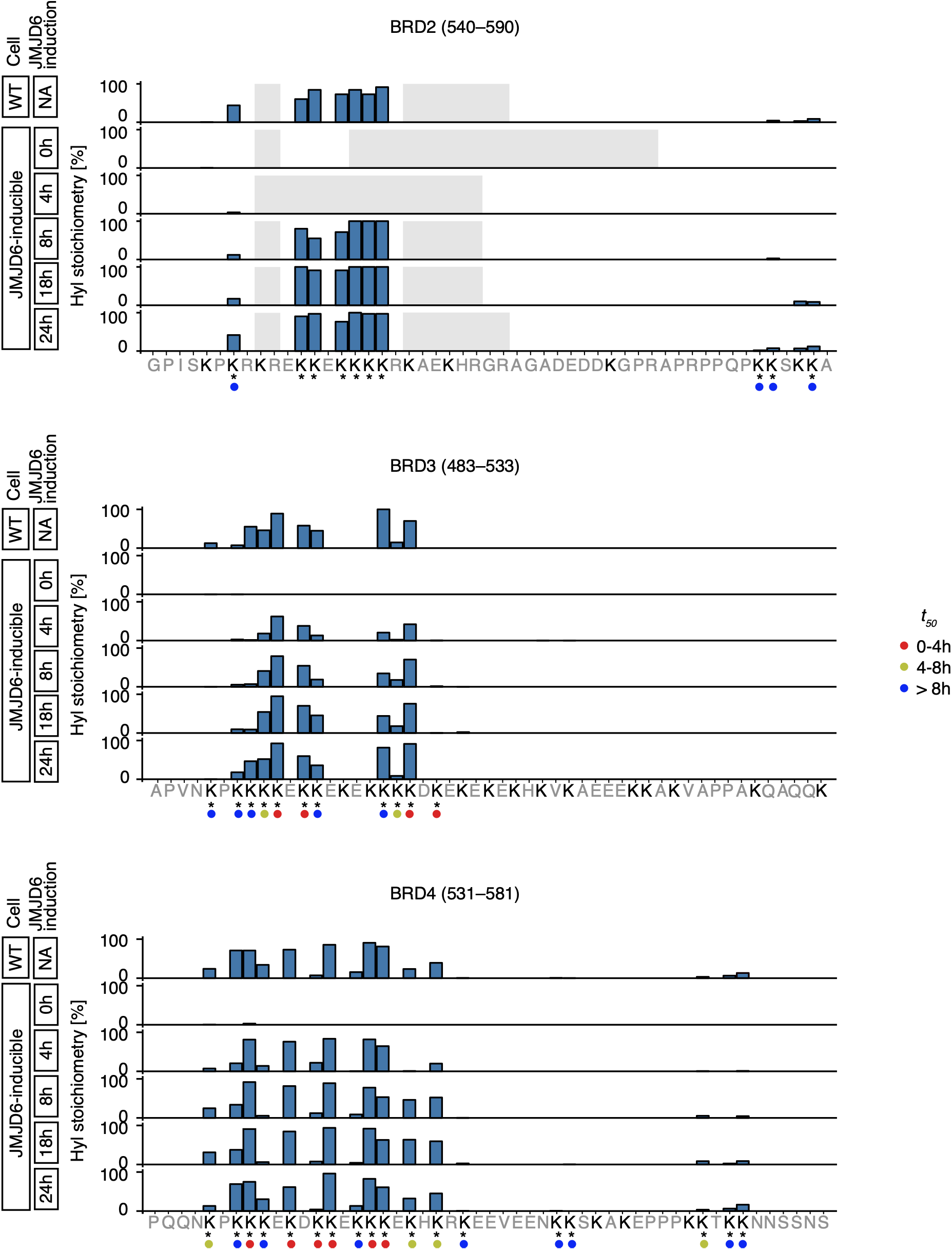
Stoichiometry of lysine hydroxylation in BRD proteins. Hydroxylysine (Hyl) stoichiometry in the lysine-rich regions of BRD2–4. Data are shown for wild-type (WT) HeLa cells and JMJD6-inducible HeLa cells following increasing durations of JMJD6 re-expression. Dots below each residue indicate its t50 category. Asterisks indicate residues with hydroxylation assignments supported by diagnostic immonium ions or assigned in our previous work^5^.

**Supplementary Fig. 7.**
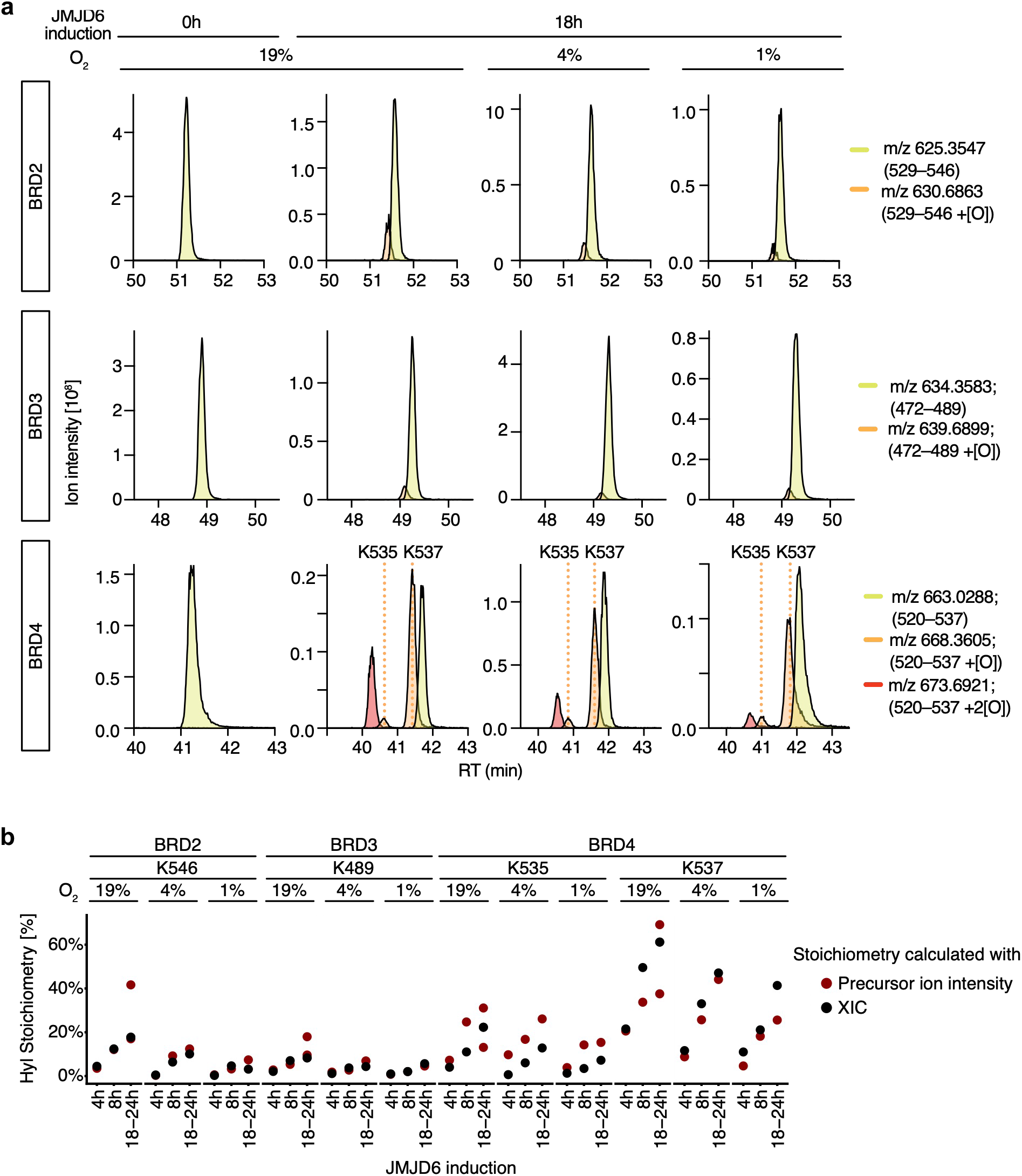
Consistent hydroxylysine stoichiometry quantification by XIC- and peptide precursor ion intensity-based methods. **a.** Extracted ion chromatograms (XICs) of the unmodified (yellow), singly hydroxylated (orange), and doubly hydroxylated (red) triply charged BRD2 (529–546), BRD3 (472–489), and BRD4 (520–537) peptides. JMJD6-inducible HeLa cells were cultured with doxycycline at 19%, 4%, or 1% O_2_ for 18h, or without doxycycline at 19% O_2_. The singly hydroxylated BRD4 peptide forms containing hydroxylation at K535 or K537 were resolved as distinct chromatographic peaks (vertical dotted lines), using a 2h gradient. **b.** Scatter plots comparing hydroxylysine (Hyl) stoichiometries at specific residues in BRD2–4, calculated from XIC analysis of non-derivatised samples (black) and from software-assigned peptide precursor ion intensities of derivatised samples (red). Data comprise 48 data points in total (4 residues × 3 oxygen levels × 3 induction durations, with independent duplicate measurements for the 18–24h time point in derivatised samples). **a** and **b**: Only residues with sufficient data coverage in non-derivatised datasets were analysed.

**Supplementary Fig. 8.**
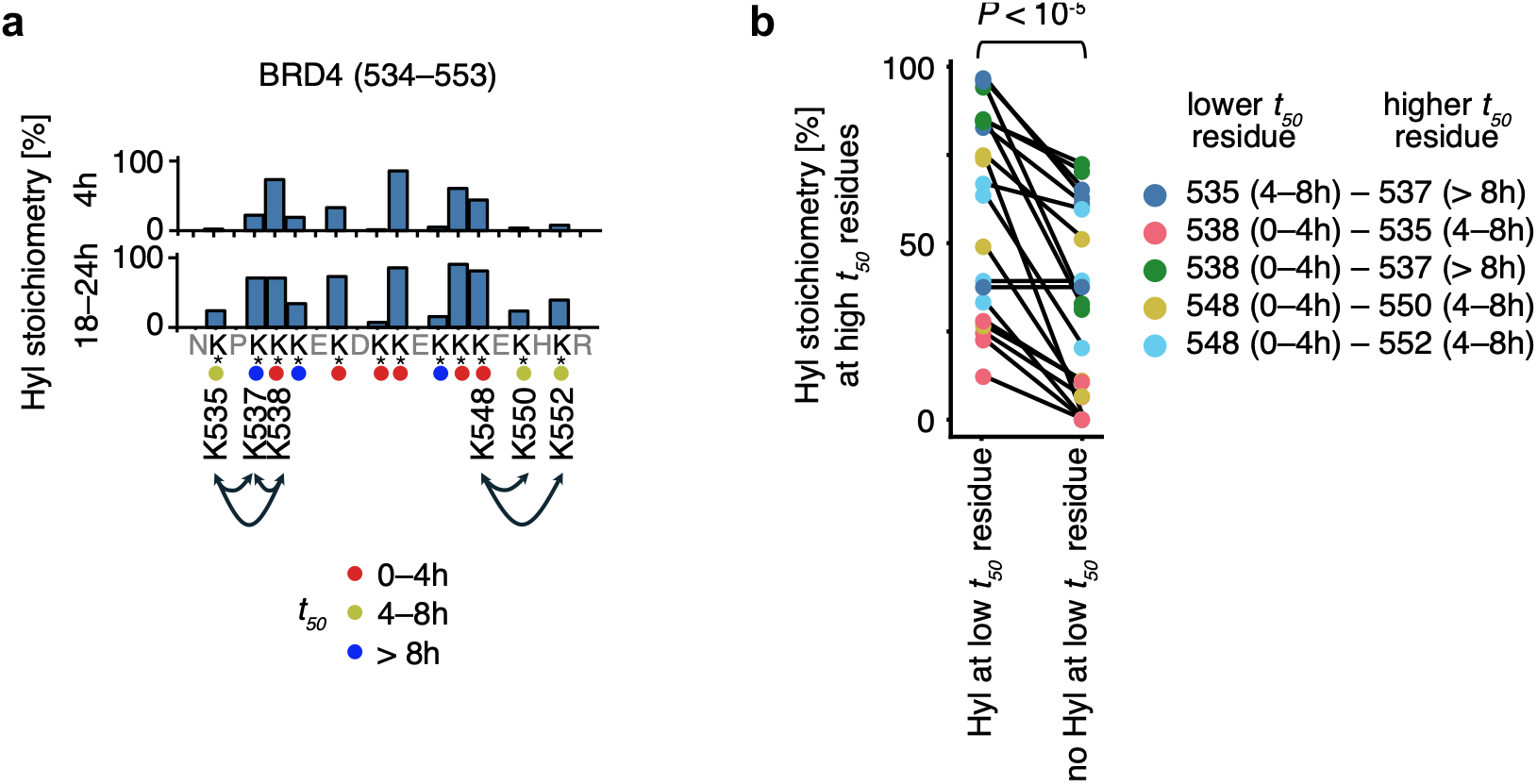
Lysine hydroxylation exhibits interdependence between neighbouring residues. **a.** Arrows show neighbouring lysine residue pairs in the BRD4 lysine-rich region (residues 534–553), spanning different *t_50_* categories, whose interdependence was analysed in **b**. For reference, hydroxylysine (Hyl) stoichiometry per residue is shown for JMJD6-inducible HeLa cells after 4 or 18–24h of JMJD6 re-expression; dots below each residue indicate its *t_50_* category. Asterisks indicate residues with hydroxylation assignments supported by diagnostic immonium ions or assigned in our previous work^5^. **b.** Hyl stoichiometry calculated from peptides in which low-*t_50_* residues were hydroxylated, compared with peptides in which the low-*t_50_* residues were not hydroxylated. The data from three independent experiments across five residue pairs were compared using a two-sided paired *t*-test.

**Supplementary Fig. 9.**
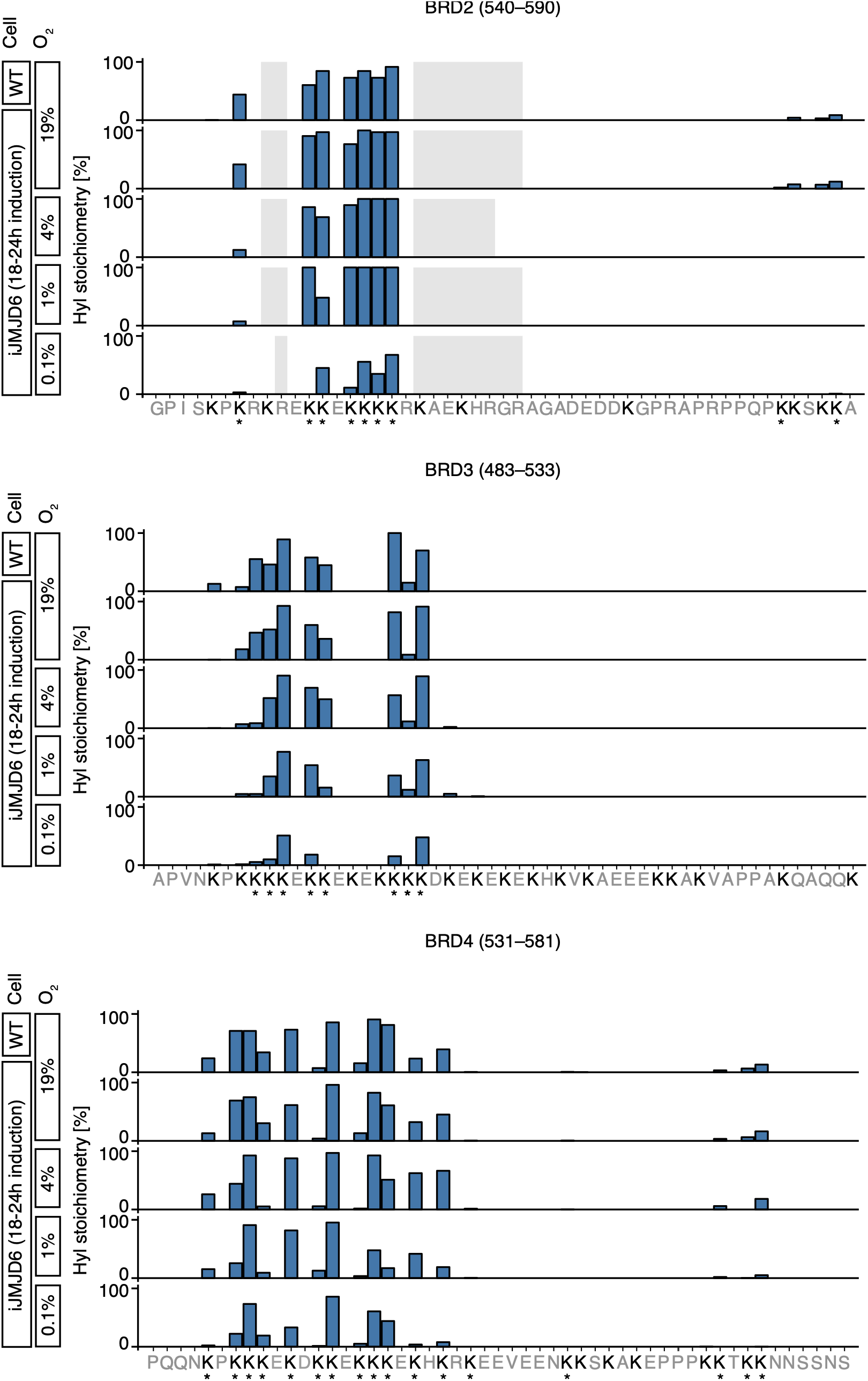
Hypoxic inhibition of lysine hydroxylation in BRD proteins. Hydroxylysine (Hyl) stoichiometry in lysine-rich regions of BRD2–4. Wild-type (WT) HeLa cells were compared with JMJD6-inducible (iJMJD6) HeLa cells following 18–24h of JMJD6 re-expression under the indicated oxygen levels. Asterisks indicate residues with hydroxylation assignments supported by diagnostic immonium ions or assigned in our previous work^5^.

**Supplementary Fig. 10.**
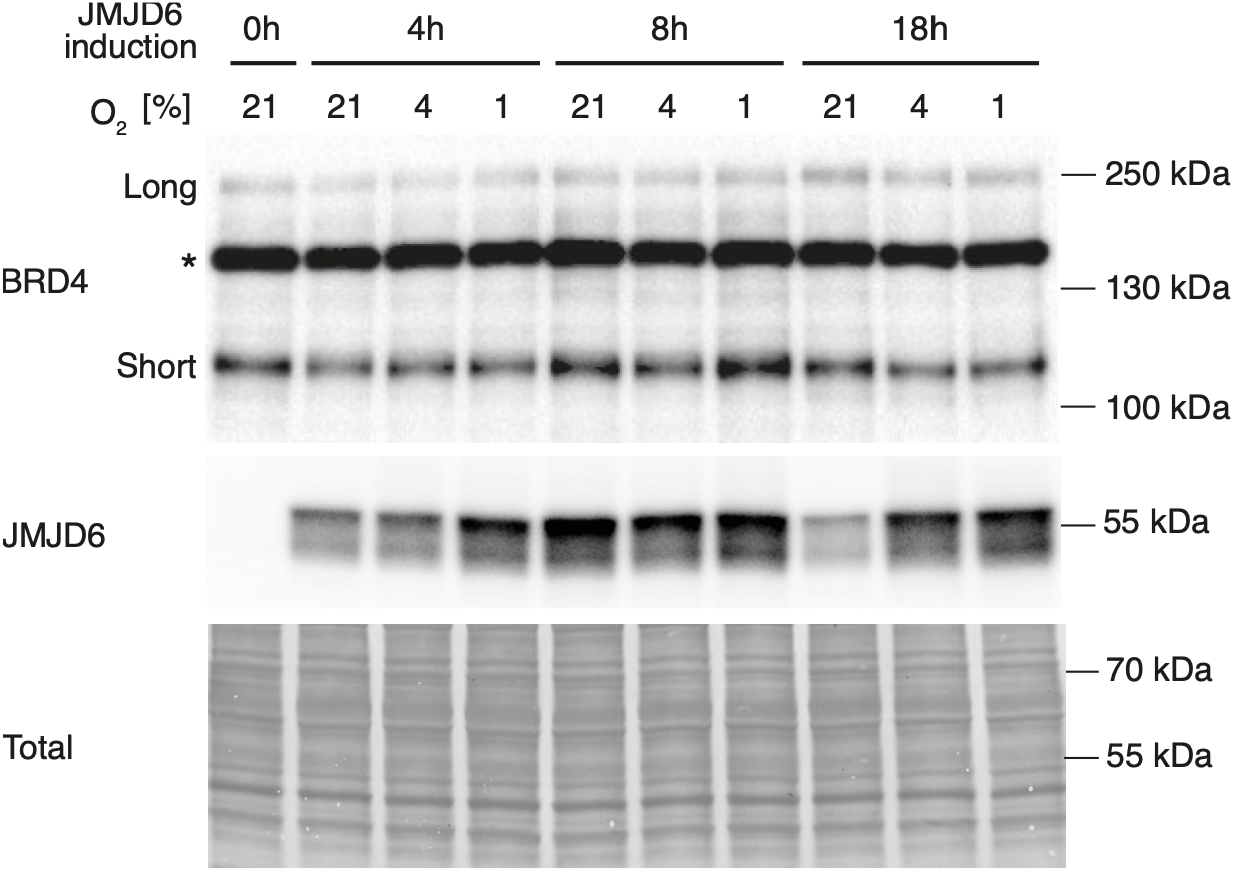
Effect of hypoxia on JMJD6 and BRD4 abundance. Immunoblotting analysis of JMJD6-inducible HeLa cells following varying durations of JMJD6 induction under different oxygen levels. Cells were treated with doxycycline for 4, 8, or 18h at 19%, 4%, or 1% O_2_, or cultured without doxycycline at 19% O_2_. The long and short isoforms of BRD4 were detected, consistent with previous studies. Non-specific species are indicated by an asterisk. Equivalent protein loading was confirmed by Coomassie staining of total protein.

**Supplementary Fig. 11.**
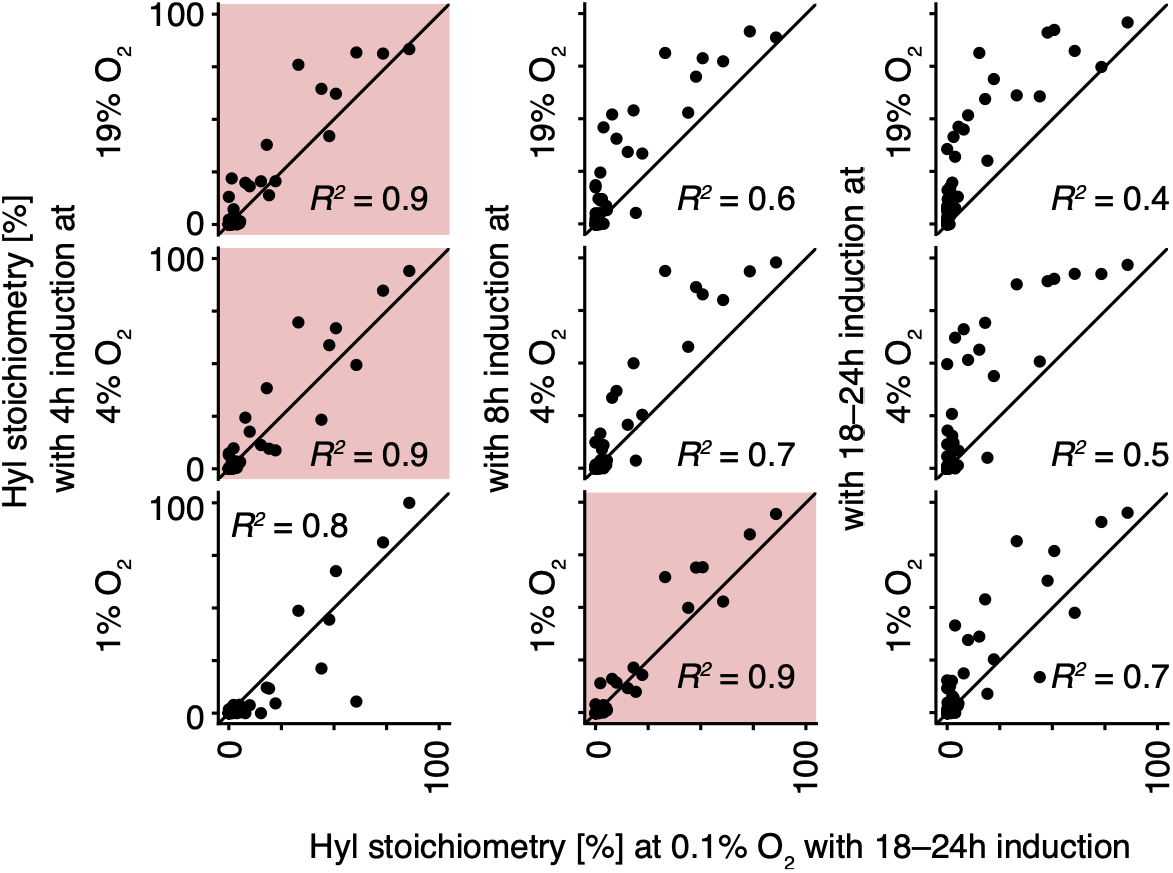
Hypoxia slows the apparent kinetics of JMJD6-catalysed lysine hydroxylation. Scatter plots comparing hydroxylysine (Hyl) stoichiometry in JMJD6-inducible HeLa cells after 18–24h of JMJD6 re-expression at 0.1% O_2_ with that measured after the indicated duration of JMJD6 re-expression at higher oxygen concentrations. For each oxygen concentration, the condition with the highest *R*^2^ value is highlighted in red.

