## Appendix for "Quantitative profiling of JMJD6-catalysed lysine hydroxylation reveals a graded, residue-dependent readout of oxygen availability"

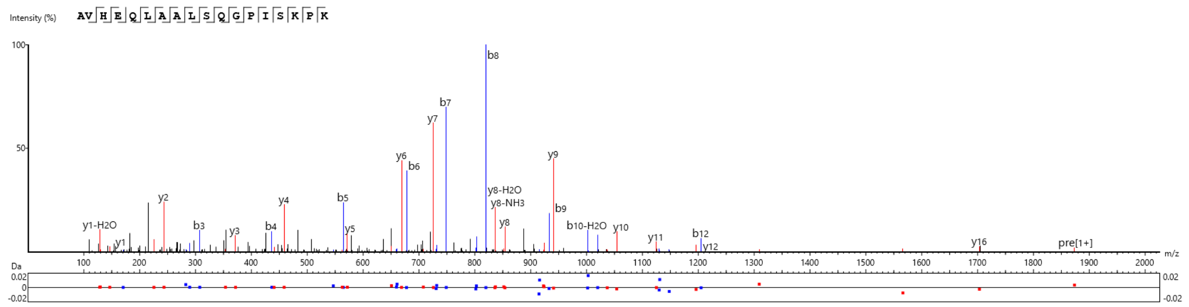


| # | b | b-H2O | b-NH3 | b(2+) | Seq | y | y-H2O | y-NH3 | y(2+) | # |
|---|---|---|---|---|---|---|---|---|---|---|
| 1 | 72.04 | 54.03 | 55.02 | 36.52 | A | | | | | 18 |
| 2 | 171.11 | 153.10 | 154.09 | 86.06 | V | 1803.01 | 1785.00 | 1785.99 | 902.01 | 17 |
| 3 | 308.17 | 290.16 | 291.14 | 154.59 | H | 1703.94 | 1685.93 | 1686.92 | 852.47 | 16 |
| 4 | 437.21 | 419.20 | 420.19 | 219.11 | E | 1566.88 | 1548.87 | 1549.86 | 783.94 | 15 |
| 5 | 565.27 | 547.26 | 548.25 | 283.14 | Q | 1437.84 | 1419.83 | 1420.82 | 719.42 | 14 |
| 6 | 678.36 | 660.35 | 661.33 | 339.68 | L | 1309.79 | 1291.77 | 1292.76 | 655.39 | 13 |
| 7 | 749.39 | 731.38 | 732.37 | 375.20 | A | 1196.70 | 1178.69 | 1179.67 | 598.85 | 12 |
| 8 | 820.43 | 802.42 | 803.41 | 410.72 | A | 1125.66 | 1107.65 | 1108.64 | 563.33 | 11 |
| 9 | 933.51 | 915.50 | 916.50 | 467.26 | L | 1054.62 | 1036.61 | 1037.60 | 527.81 | 10 |
| 10 | 1020.55 | 1002.54 | 1003.54 | 510.77 | S | 941.54 | 923.53 | 924.52 | 471.27 | 9 |
| 11 | 1148.60 | 1130.59 | 1131.59 | 574.80 | Q | 854.51 | 836.50 | 837.48 | 427.75 | 8 |
| 12 | 1205.63 | 1187.62 | 1188.60 | 603.31 | G | 726.45 | 708.44 | 709.42 | 363.73 | 7 |
| 13 | 1302.68 | 1284.67 | 1285.65 | 651.84 | P | 669.43 | 651.42 | 652.40 | 335.21 | 6 |
| 14 | 1415.76 | 1397.75 | 1398.74 | 708.38 | I | 572.38 | 554.37 | 555.35 | 286.69 | 5 |
| 15 | 1502.80 | 1484.79 | 1485.77 | 751.90 | S | 459.29 | 441.28 | 442.27 | 230.15 | 4 |
| 16 | 1630.89 | 1612.88 | 1613.86 | 815.95 | K | 372.26 | 354.25 | 355.23 | 186.63 | 3 |
| 17 | 1727.94 | 1709.93 | 1710.92 | 864.47 | P | 244.17 | 226.15 | 227.14 | 122.58 | 2 |
| 18 | | | | | K | 147.11 | 129.10 | 130.09 | 74.06 | 1 |

### Representative MS/MS spectrum of the unmodified BRD2(529–546) peptide

MS/MS spectrum and ion table of the unmodified BRD2(529–546) peptide in uninduced JMJD6-inducible HeLa cells at 19% $O_2$; scan 19353; $P < 10^{-40}$.

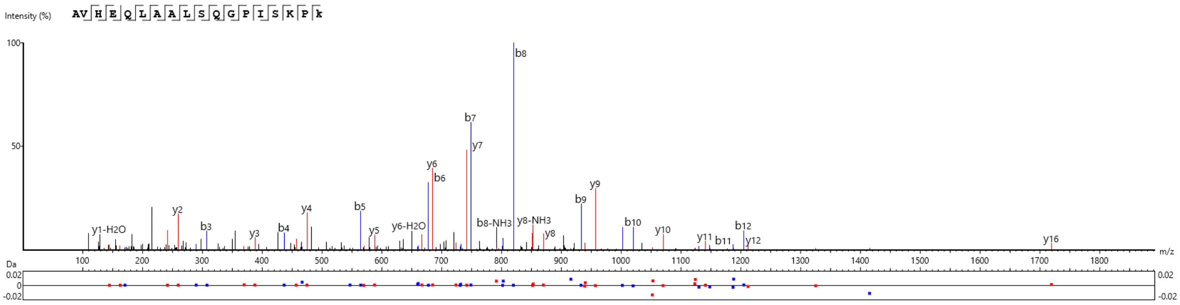


| # | b | b-H2O | b-NH3 | b(2+) | Seq | y | y-H2O | y-NH3 | y(2+) | # |
|---|---|---|---|---|---|---|---|---|---|---|
| 1 | 72.04 | 54.03 | 55.02 | 36.52 | A | | | | | 18 |
| 2 | 171.11 | 153.10 | 154.09 | 86.06 | V | 1819.01 | 1801.00 | 1801.98 | 910.00 | 17 |
| 3 | 308.17 | 290.16 | 291.14 | 154.59 | H | 1719.94 | 1701.93 | 1702.91 | 860.47 | 16 |
| 4 | 437.21 | 419.20 | 420.19 | 219.11 | E | 1582.88 | 1564.87 | 1565.85 | 791.95 | 15 |
| 5 | 565.27 | 547.26 | 548.25 | 283.14 | Q | 1453.84 | 1435.83 | 1436.81 | 727.42 | 14 |
| 6 | 678.36 | 660.35 | 661.33 | 339.68 | L | 1325.78 | 1307.77 | 1308.75 | 663.39 | 13 |
| 7 | 749.39 | 731.38 | 732.37 | 375.20 | A | 1212.69 | 1194.68 | 1195.67 | 606.85 | 12 |
| 8 | 820.43 | 802.42 | 803.41 | 410.72 | A | 1141.66 | 1123.65 | 1124.64 | 571.33 | 11 |
| 9 | 933.52 | 915.50 | 916.50 | 467.26 | L | 1070.62 | 1052.60 | 1053.60 | 535.81 | 10 |
| 10 | 1020.55 | 1002.54 | 1003.52 | 510.77 | S | 957.54 | 939.52 | 940.51 | 479.27 | 9 |
| 11 | 1148.60 | 1130.59 | 1131.58 | 574.80 | Q | 870.50 | 852.49 | 853.48 | 435.75 | 8 |
| 12 | 1205.63 | 1187.61 | 1188.61 | 603.31 | G | 742.45 | 724.44 | 725.42 | 371.72 | 7 |
| 13 | 1302.68 | 1284.67 | 1285.65 | 651.84 | P | 685.42 | 667.41 | 668.40 | 343.21 | 6 |
| 14 | 1415.75 | 1397.75 | 1398.74 | 708.38 | I | 588.37 | 570.36 | 571.34 | 294.69 | 5 |
| 15 | 1502.80 | 1484.79 | 1485.77 | 751.90 | S | 475.29 | 457.28 | 458.26 | 238.14 | 4 |
| 16 | 1630.89 | 1612.88 | 1613.86 | 815.95 | K | 388.26 | 370.25 | 371.23 | 194.63 | 3 |
| 17 | 1727.94 | 1709.93 | 1710.92 | 864.47 | P | 260.16 | 242.15 | 243.13 | 130.58 | 2 |
| 18 | | | | | K(+15.99) | 163.11 | 145.10 | 146.08 | 82.05 | 1 |

**Representative MS/MS spectrum of the K546-hydroxylated BRD2(529–546) peptide**

MS/MS spectrum and ion table of the K546-hydroxylated BRD2(529–546) peptide in JMJD6-inducible HeLa cells following incubation with doxycycline for 18h at 19% $O_2$; scan 19303; $P < 10^{-40}$.

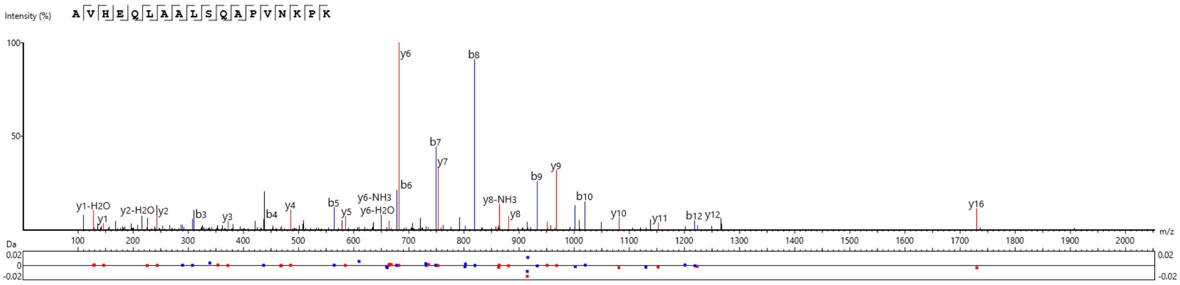


| # | b | b-H2O | b-NH3 | b(2+) | Seq | y | y-H2O | y-NH3 | y(2+) | # |
|---|---|---|---|---|---|---|---|---|---|---|
| 1 | 72.04 | 54.03 | 55.02 | 36.52 | A | | | | | 18 |
| 2 | 171.11 | 153.10 | 154.09 | 86.06 | V | 1830.02 | 1812.01 | 1813.00 | 915.50 | 17 |
| 3 | 308.17 | 290.16 | 291.14 | 154.59 | H | 1730.95 | 1712.94 | 1713.93 | 865.98 | 16 |
| 4 | 437.21 | 419.20 | 420.19 | 219.11 | E | 1593.90 | 1575.89 | 1576.87 | 797.45 | 15 |
| 5 | 565.27 | 547.26 | 548.25 | 283.14 | Q | 1464.85 | 1446.84 | 1447.83 | 732.93 | 14 |
| 6 | 678.36 | 660.34 | 661.33 | 339.68 | L | 1336.79 | 1318.78 | 1319.77 | 668.90 | 13 |
| 7 | 749.39 | 731.39 | 732.37 | 375.20 | A | 1223.71 | 1205.70 | 1206.68 | 612.36 | 12 |
| 8 | 820.43 | 802.42 | 803.41 | 410.72 | A | 1152.67 | 1134.66 | 1135.65 | 576.84 | 11 |
| 9 | 933.51 | 915.50 | 916.50 | 467.26 | L | 1081.63 | 1063.63 | 1064.61 | 541.32 | 10 |
| 10 | 1020.55 | 1002.53 | 1003.52 | 510.77 | S | 968.55 | 950.54 | 951.53 | 484.78 | 9 |
| 11 | 1148.61 | 1130.59 | 1131.58 | 574.80 | Q | 881.52 | 863.51 | 864.49 | 441.26 | 8 |
| 12 | 1219.64 | 1201.63 | 1202.62 | 610.33 | A | 753.46 | 735.45 | 736.44 | 377.23 | 7 |
| 13 | 1316.70 | 1298.69 | 1299.67 | 658.85 | P | 682.42 | 664.41 | 665.40 | 341.71 | 6 |
| 14 | 1415.76 | 1397.75 | 1398.74 | 708.38 | V | 585.37 | 567.36 | 568.34 | 293.19 | 5 |
| 15 | 1529.81 | 1511.80 | 1512.78 | 765.40 | N | 486.30 | 468.29 | 469.28 | 243.65 | 4 |
| 16 | 1657.90 | 1639.89 | 1640.88 | 829.45 | K | 372.26 | 354.25 | 355.23 | 186.63 | 3 |
| 17 | 1754.95 | 1736.94 | 1737.93 | 877.98 | P | 244.17 | 226.15 | 227.14 | 122.58 | 2 |
| 18 | | | | | K | 147.11 | 129.10 | 130.09 | 74.06 | 1 |

**Representative MS/MS spectrum of the unmodified BRD3(472–489) peptide**

MS/MS spectrum and ion table of the unmodified BRD3(472–489) peptide in uninduced JMJD6-inducible HeLa cells at 19% $O_2$; scan 18213; $P < 10^{-40}$.

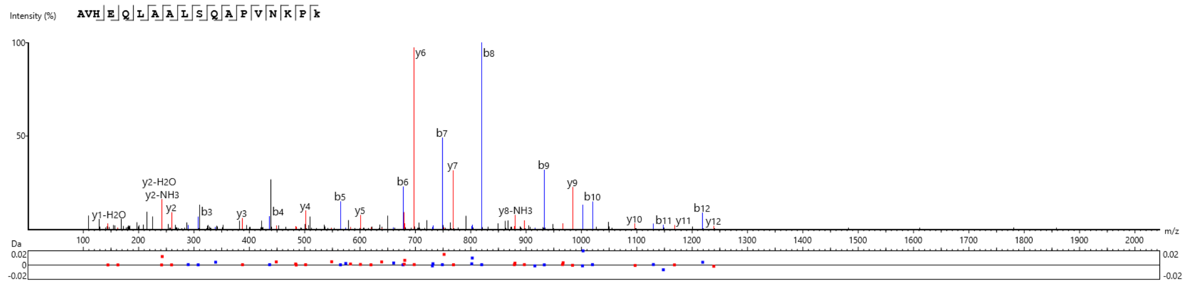


| # | b | b-H2O | b-NH3 | b(2+) | Seq | y | y-H2O | y-NH3 | y(2+) | # |
|---|---|---|---|---|---|---|---|---|---|---|
| 1 | 72.04 | 54.03 | 55.02 | 36.52 | A | | | | | 18 |
| 2 | 171.11 | 153.10 | 154.09 | 86.06 | V | 1846.02 | 1828.01 | 1828.99 | 923.51 | 17 |
| 3 | 308.17 | 290.16 | 291.14 | 154.59 | H | 1746.95 | 1728.94 | 1729.92 | 873.97 | 16 |
| 4 | 437.21 | 419.20 | 420.19 | 219.11 | E | 1609.89 | 1591.88 | 1592.86 | 805.45 | 15 |
| 5 | 565.27 | 547.26 | 548.25 | 283.14 | Q | 1480.85 | 1462.84 | 1463.82 | 740.92 | 14 |
| 6 | 678.36 | 660.35 | 661.33 | 339.68 | L | 1352.79 | 1334.78 | 1335.76 | 676.89 | 13 |
| 7 | 749.39 | 731.38 | 732.37 | 375.20 | A | 1239.70 | 1221.69 | 1222.68 | 620.35 | 12 |
| 8 | 820.43 | 802.42 | 803.41 | 410.72 | A | 1168.67 | 1150.66 | 1151.64 | 584.83 | 11 |
| 9 | 933.52 | 915.50 | 916.49 | 467.26 | L | 1097.63 | 1079.62 | 1080.60 | 549.32 | 10 |
| 10 | 1020.55 | 1002.54 | 1003.54 | 510.77 | S | 984.55 | 966.54 | 967.52 | 492.77 | 9 |
| 11 | 1148.60 | 1130.60 | 1131.58 | 574.80 | Q | 897.52 | 879.50 | 880.49 | 449.26 | 8 |
| 12 | 1219.65 | 1201.63 | 1202.62 | 610.32 | A | 769.46 | 751.45 | 752.44 | 385.23 | 7 |
| 13 | 1316.70 | 1298.69 | 1299.67 | 658.85 | P | 698.42 | 680.41 | 681.40 | 349.71 | 6 |
| 14 | 1415.76 | 1397.75 | 1398.74 | 708.38 | V | 601.37 | 583.36 | 584.34 | 301.18 | 5 |
| 15 | 1529.81 | 1511.80 | 1512.78 | 765.40 | N | 502.30 | 484.29 | 485.27 | 251.65 | 4 |
| 16 | 1657.90 | 1639.89 | 1640.88 | 829.45 | K | 388.26 | 370.24 | 371.23 | 194.63 | 3 |
| 17 | 1754.95 | 1736.94 | 1737.93 | 877.98 | P | 260.16 | 242.15 | 243.14 | 130.58 | 2 |
| 18 | | | | | K(+15.99) | 163.11 | 145.10 | 146.08 | 82.05 | 1 |

**Representative MS/MS spectrum of the K489-hydroxylated BRD3(472–489) peptide**

MS/MS spectrum and ion table of the K489-hydroxylated BRD3(472–489) peptide in JMJD6-inducible HeLa cells following incubation with doxycycline for 18h at 19% $O_2$; scan 17893; $P < 10^{-40}$.

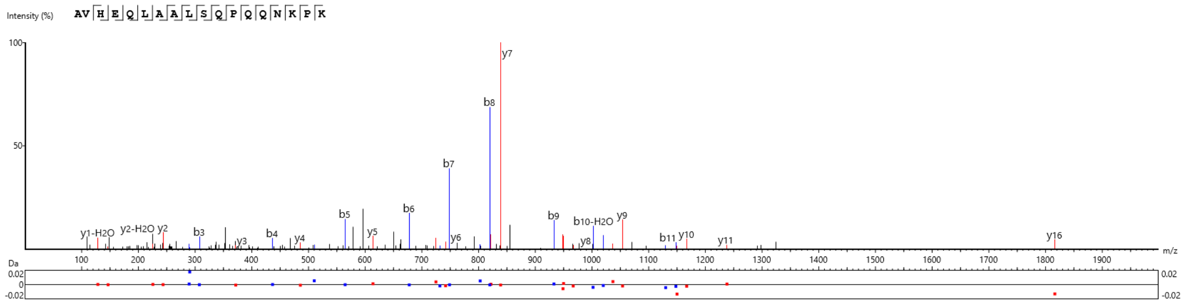


| # | b | b-H2O | b-NH3 | b(2+) | Seq | y | y-H2O | y-NH3 | y(2+) | # |
|---|---|---|---|---|---|---|---|---|---|---|
| 1 | 72.04 | 54.03 | 55.02 | 36.52 | A | | | | | 18 |
| 2 | 171.11 | 153.10 | 154.09 | 86.06 | V | 1916.03 | 1898.02 | 1899.01 | 958.52 | 17 |
| 3 | 308.17 | 290.16 | 291.16 | 154.59 | H | 1816.95 | 1798.96 | 1799.94 | 908.98 | 16 |
| 4 | 437.21 | 419.20 | 420.19 | 219.11 | E | 1679.91 | 1661.90 | 1662.88 | 840.45 | 15 |
| 5 | 565.27 | 547.26 | 548.25 | 283.14 | Q | 1550.86 | 1532.85 | 1533.84 | 775.93 | 14 |
| 6 | 678.36 | 660.35 | 661.33 | 339.68 | L | 1422.81 | 1404.80 | 1405.78 | 711.90 | 13 |
| 7 | 749.39 | 731.38 | 732.36 | 375.20 | A | 1309.72 | 1291.71 | 1292.70 | 655.36 | 12 |
| 8 | 820.43 | 802.42 | 803.41 | 410.72 | A | 1238.69 | 1220.67 | 1221.66 | 619.84 | 11 |
| 9 | 933.52 | 915.50 | 916.49 | 467.26 | L | 1167.65 | 1149.64 | 1150.61 | 584.32 | 10 |
| 10 | 1020.55 | 1002.53 | 1003.52 | 510.78 | S | 1054.56 | 1036.55 | 1037.54 | 527.78 | 9 |
| 11 | 1148.60 | 1130.59 | 1131.58 | 574.80 | Q | 967.53 | 949.52 | 950.51 | 484.27 | 8 |
| 12 | 1245.66 | 1227.65 | 1228.63 | 623.33 | P | 839.47 | 821.46 | 822.45 | 420.24 | 7 |
| 13 | 1373.72 | 1355.71 | 1356.69 | 687.36 | Q | 742.42 | 724.41 | 725.40 | 371.71 | 6 |
| 14 | 1501.78 | 1483.77 | 1484.75 | 751.39 | Q | 614.36 | 596.35 | 597.34 | 307.68 | 5 |
| 15 | 1615.82 | 1597.81 | 1598.79 | 808.41 | N | 486.30 | 468.29 | 469.28 | 243.65 | 4 |
| 16 | 1743.91 | 1725.90 | 1726.89 | 872.46 | K | 372.26 | 354.25 | 355.23 | 186.63 | 3 |
| 17 | 1840.97 | 1822.96 | 1823.94 | 920.98 | P | 244.16 | 226.15 | 227.14 | 122.58 | 2 |
| 18 | | | | | K | 147.11 | 129.10 | 130.09 | 74.06 | 1 |

**Representative MS/MS spectrum of the unmodified BRD4(520–537) peptide**

MS/MS spectrum and ion table of the unmodified BRD4(520–537) peptide in uninduced JMJD6-inducible HeLa cells at 19% $O_2$; scan 13877; $P < 10^{-40}$.

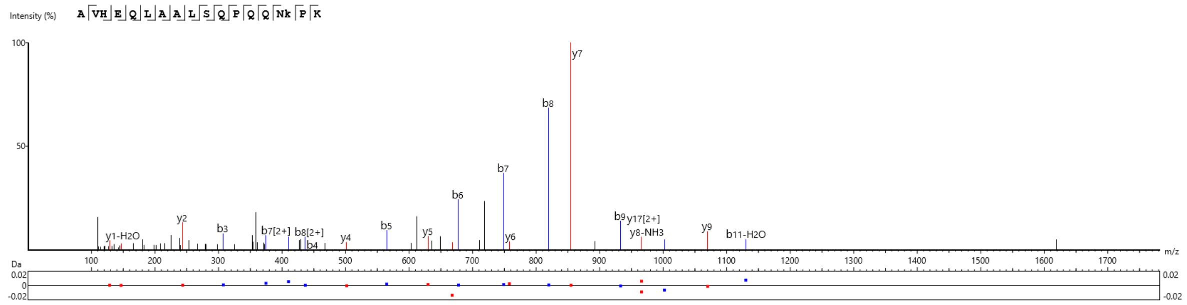


| # | b | b-H2O | b-NH3 | b(2+) | Seq | y | y-H2O | y-NH3 | y(2+) | # |
|---|---|---|---|---|---|---|---|---|---|---|
| 1 | 72.04 | 54.03 | 55.02 | 36.52 | A | | | | | 18 |
| 2 | 171.11 | 153.10 | 154.09 | 86.06 | V | 1932.03 | 1914.02 | 1915.00 | 966.51 | 17 |
| 3 | 308.17 | 290.16 | 291.14 | 154.59 | H | 1832.96 | 1814.95 | 1815.93 | 916.98 | 16 |
| 4 | 437.21 | 419.20 | 420.19 | 219.11 | E | 1695.90 | 1677.89 | 1678.88 | 848.45 | 15 |
| 5 | 565.27 | 547.26 | 548.25 | 283.14 | Q | 1566.86 | 1548.85 | 1549.83 | 783.93 | 14 |
| 6 | 678.36 | 660.35 | 661.33 | 339.68 | L | 1438.80 | 1420.79 | 1421.77 | 719.90 | 13 |
| 7 | 749.39 | 731.38 | 732.37 | 375.20 | A | 1325.72 | 1307.71 | 1308.69 | 663.36 | 12 |
| 8 | 820.43 | 802.42 | 803.40 | 410.72 | A | 1254.68 | 1236.67 | 1237.65 | 627.84 | 11 |
| 9 | 933.51 | 915.50 | 916.49 | 467.26 | L | 1183.64 | 1165.63 | 1166.62 | 592.32 | 10 |
| 10 | 1020.55 | 1002.53 | 1003.52 | 510.77 | S | 1070.56 | 1052.55 | 1053.53 | 535.78 | 9 |
| 11 | 1148.61 | 1130.60 | 1131.58 | 574.80 | Q | 983.53 | 965.52 | 966.51 | 492.26 | 8 |
| 12 | 1245.66 | 1227.65 | 1228.63 | 623.33 | P | 855.47 | 837.46 | 838.44 | 428.23 | 7 |
| 13 | 1373.72 | 1355.71 | 1356.69 | 687.36 | Q | 758.42 | 740.40 | 741.39 | 379.71 | 6 |
| 14 | 1501.78 | 1483.77 | 1484.75 | 751.39 | Q | 630.36 | 612.35 | 613.33 | 315.68 | 5 |
| 15 | 1615.82 | 1597.81 | 1598.79 | 808.41 | N | 502.30 | 484.29 | 485.27 | 251.65 | 4 |
| 16 | 1759.91 | 1741.90 | 1742.88 | 880.45 | K(+15.99) | 388.26 | 370.24 | 371.23 | 194.63 | 3 |
| 17 | 1856.96 | 1838.95 | 1839.93 | 928.98 | P | 244.17 | 226.15 | 227.14 | 122.58 | 2 |
| 18 | | | | | K | 147.11 | 129.10 | 130.09 | 74.06 | 1 |

**Representative MS/MS spectrum of the K535-hydroxylated BRD4(520–537) peptide**

MS/MS spectrum and ion table of the K535-hydroxylated BRD4(520–537) peptide in JMJD6-inducible HeLa cells following incubation with doxycycline for 18h at 19% $O_2$; scan 13382; $P < 10^{-40}$.

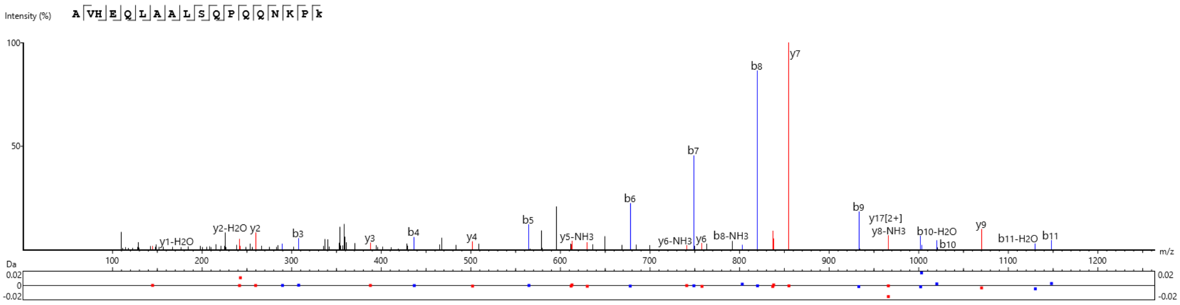


| # | b | b-H2O | b-NH3 | b(2+) | Seq | y | y-H2O | y-NH3 | y(2+) | # |
|---|---|---|---|---|---|---|---|---|---|---|
| 1 | 72.04 | 54.03 | 55.02 | 36.52 | A | | | | | 18 |
| 2 | 171.11 | 153.10 | 154.09 | 86.06 | V | 1932.03 | 1914.02 | 1915.00 | 966.50 | 17 |
| 3 | 308.17 | 290.16 | 291.14 | 154.59 | H | 1832.96 | 1814.95 | 1815.93 | 916.98 | 16 |
| 4 | 437.21 | 419.20 | 420.19 | 219.11 | E | 1695.90 | 1677.89 | 1678.88 | 848.45 | 15 |
| 5 | 565.27 | 547.26 | 548.25 | 283.14 | Q | 1566.86 | 1548.85 | 1549.83 | 783.93 | 14 |
| 6 | 678.36 | 660.35 | 661.33 | 339.68 | L | 1438.80 | 1420.79 | 1421.77 | 719.90 | 13 |
| 7 | 749.39 | 731.38 | 732.37 | 375.20 | A | 1325.72 | 1307.71 | 1308.69 | 663.36 | 12 |
| 8 | 820.43 | 802.42 | 803.41 | 410.72 | A | 1254.68 | 1236.67 | 1237.65 | 627.84 | 11 |
| 9 | 933.51 | 915.50 | 916.49 | 467.26 | L | 1183.64 | 1165.63 | 1166.62 | 592.32 | 10 |
| 10 | 1020.55 | 1002.53 | 1003.54 | 510.77 | S | 1070.56 | 1052.55 | 1053.53 | 535.78 | 9 |
| 11 | 1148.61 | 1130.59 | 1131.58 | 574.80 | Q | 983.53 | 965.52 | 966.50 | 492.26 | 8 |
| 12 | 1245.66 | 1227.65 | 1228.63 | 623.33 | P | 855.47 | 837.46 | 838.44 | 428.23 | 7 |
| 13 | 1373.72 | 1355.71 | 1356.69 | 687.36 | Q | 758.41 | 740.40 | 741.39 | 379.71 | 6 |
| 14 | 1501.78 | 1483.77 | 1484.75 | 751.39 | Q | 630.36 | 612.35 | 613.33 | 315.68 | 5 |
| 15 | 1615.82 | 1597.81 | 1598.79 | 808.41 | N | 502.30 | 484.29 | 485.27 | 251.65 | 4 |
| 16 | 1743.91 | 1725.90 | 1726.89 | 872.46 | K | 388.26 | 370.24 | 371.23 | 194.63 | 3 |
| 17 | 1840.97 | 1822.96 | 1823.94 | 920.98 | P | 260.16 | 242.15 | 243.14 | 130.58 | 2 |
| 18 | | | | | K(+15.99) | 163.11 | 145.10 | 146.08 | 82.05 | 1 |

**Representative MS/MS spectrum of the K537-hydroxylated BRD4(520–537) peptide**

MS/MS spectrum and ion table of the K537-hydroxylated BRD4(520–537) peptide in JMJD6-inducible HeLa cells following incubation with doxycycline for 18h at 19% $O_2$; scan 13726; $P < 10^{-40}$.

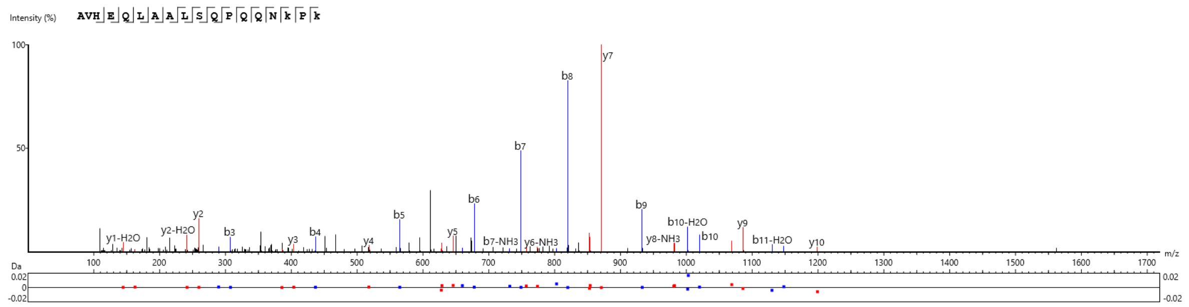


| # | b | b-H2O | b-NH3 | b(2+) | Seq | y | y-H2O | y-NH3 | y(2+) | # |
|---|---|---|---|---|---|---|---|---|---|---|
| 1 | 72.04 | 54.03 | 55.02 | 36.52 | A | | | | | 18 |
| 2 | 171.11 | 153.10 | 154.09 | 86.06 | V | 1948.02 | 1930.01 | 1931.00 | 974.51 | 17 |
| 3 | 308.17 | 290.16 | 291.14 | 154.59 | H | 1848.96 | 1830.95 | 1831.93 | 924.98 | 16 |
| 4 | 437.21 | 419.20 | 420.19 | 219.11 | E | 1711.90 | 1693.89 | 1694.87 | 856.45 | 15 |
| 5 | 565.27 | 547.26 | 548.25 | 283.14 | Q | 1582.85 | 1564.84 | 1565.83 | 791.93 | 14 |
| 6 | 678.36 | 660.35 | 661.33 | 339.68 | L | 1454.80 | 1436.79 | 1437.77 | 727.90 | 13 |
| 7 | 749.39 | 731.38 | 732.37 | 375.20 | A | 1341.71 | 1323.70 | 1324.69 | 671.36 | 12 |
| 8 | 820.43 | 802.42 | 803.41 | 410.72 | A | 1270.67 | 1252.66 | 1253.65 | 635.84 | 11 |
| 9 | 933.51 | 915.50 | 916.49 | 467.26 | L | 1199.63 | 1181.63 | 1182.61 | 600.32 | 10 |
| 10 | 1020.55 | 1002.53 | 1003.54 | 510.77 | S | 1086.55 | 1068.54 | 1069.53 | 543.78 | 9 |
| 11 | 1148.61 | 1130.59 | 1131.58 | 574.80 | Q | 999.52 | 981.51 | 982.50 | 500.26 | 8 |
| 12 | 1245.66 | 1227.65 | 1228.63 | 623.33 | P | 871.46 | 853.45 | 854.44 | 436.23 | 7 |
| 13 | 1373.72 | 1355.71 | 1356.69 | 687.36 | Q | 774.41 | 756.40 | 757.39 | 387.71 | 6 |
| 14 | 1501.78 | 1483.77 | 1484.75 | 751.39 | Q | 646.35 | 628.34 | 629.33 | 323.68 | 5 |
| 15 | 1615.82 | 1597.81 | 1598.79 | 808.41 | N | 518.29 | 500.28 | 501.27 | 259.65 | 4 |
| 16 | 1759.91 | 1741.90 | 1742.88 | 880.45 | K(+15.99) | 404.25 | 386.24 | 387.22 | 202.63 | 3 |
| 17 | 1856.96 | 1838.95 | 1839.93 | 928.98 | P | 260.16 | 242.15 | 243.13 | 130.58 | 2 |
| 18 | | | | | K(+15.99) | 163.11 | 145.10 | 146.08 | 82.05 | 1 |

**Representative MS/MS spectrum of BRD4(520–537) peptide doubly hydroxylated at K535 and K537**

MS/MS spectrum and ion table of the BRD4(520–537) peptide doubly hydroxylated at K535 and K537 in JMJD6-inducible HeLa cells following incubation with doxycycline for 18h at 19% $O_2$; scan 13229; $P < 10^{-40}$.
